# FcγRIIB regulates (auto)antibody responses by limiting marginal zone B cell activation

**DOI:** 10.1101/2021.12.03.471075

**Authors:** Ashley N. Barlev, Susan Malkiel, Annemarie L. Dorjée, Jolien Suurmond, Betty Diamond

## Abstract

FcγRIIB is an inhibitory receptor expressed throughout B cell development. Diminished expression or function is associated with lupus in mice and humans, in particular through an effect on autoantibody production and plasma cell differentiation. Here, we analysed the effect of B cell-intrinsic FcγRIIB expression on B cell activation and plasma cell differentiation.

Loss of FcγRIIB on B cells (*Fcgr2b* cKO mice) led to a spontaneous increase in autoantibody titers. This increase was most striking for IgG3, suggestive of increased extrafollicular responses. Marginal zone (MZ) and IgG3+ B cells had the highest expression of FcγRIIB and the increase in serum IgG3 was linked to increased MZ B cell signaling and activation in the absence of FcγRIIB. Likewise, human circulating MZ-like B cells had the highest expression of FcγRIIB, and their activation was most strongly inhibited by engaging FcγRIIB. Finally, marked increases in IgG3+ plasma cells and B cells were observed during extrafollicular plasma cell responses with both T-dependent and T-independent antigens in *Fcgr2b* cKO mice. The increased IgG3 response following immunization of *Fcgr2b* cKO mice was lost in MZ-deficient *Notch2*/*Fcgr2b* cKO mice.

Thus, we present a model where high FcγRIIB expression in MZ B cells prevents their hyperactivation and ensuing autoimmunity.

**Graphical abstract:** 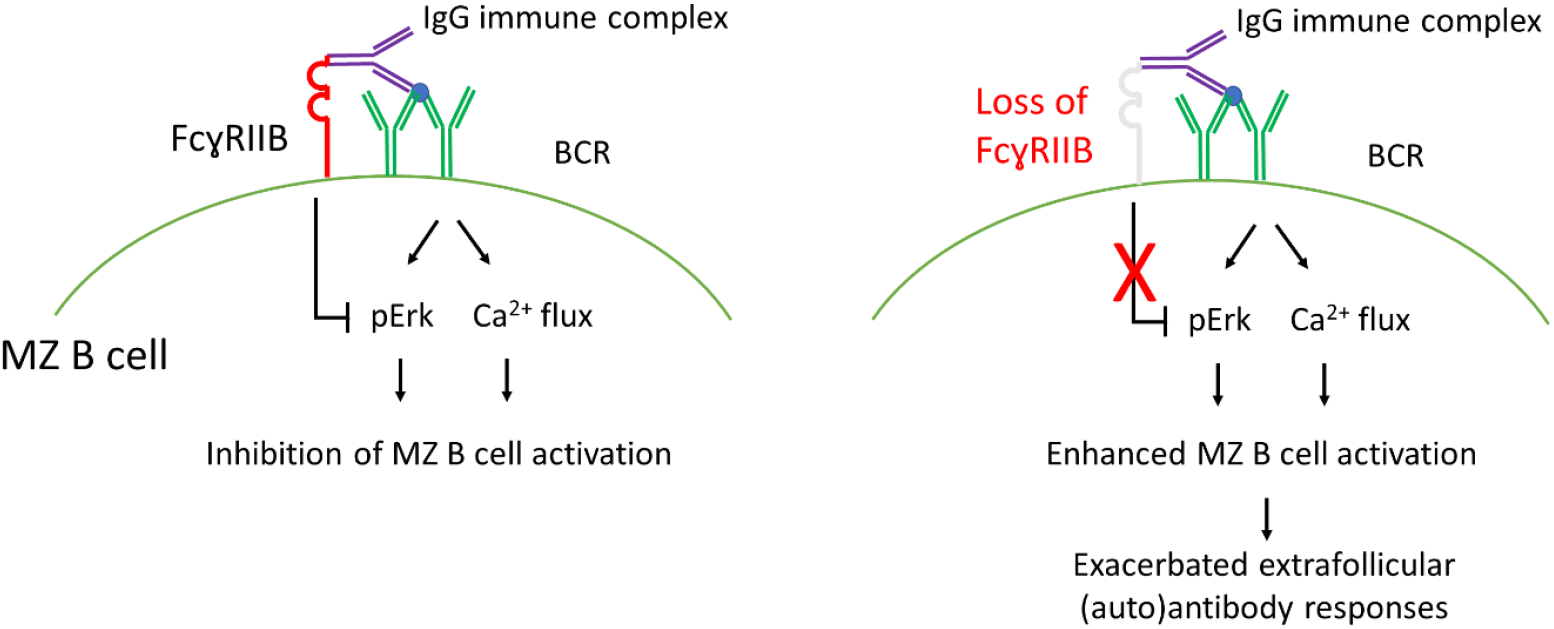

## Introduction

FcγRIIB is an inhibitory receptor expressed on many cell types, including B cells. On B cells, it is the only Fcγ receptor and is expressed throughout B cell development. When FcγRIIB is crosslinked to the B cell receptor (BCR), downstream BCR signaling and B cell activation is inhibited (1, 2). There are alleles of *FCGR2B* leading to loss of function or reduced expression that predispose to systemic lupus erythematosus (SLE) (3-6). The most common of these SNPs, the FcγRIIB -T232 variant, acts through loss of motility of FcγRIIB in the membrane preventing proximity to the BCR (1, 6-8).

In mice, global FcγRIIB deficiency was initially reported to cause a lupus-like disease, with presence of autoantibodies and deposition of immune complexes in the kidney (9). There has, however, been controversy surrounding the predisposition to lupus in FcγRIIB deficient mice, as some studies suggest that other genes in concordance with *Fcgr2b* are required to influence the development of lupus (10, 11). The effect of FcγRIIB on autoantibody production is B cell intrinsic, as overexpression of FcγRIIB on B cells from lupus-prone mice leads to improvement in lupus phenotype (12, 13) and B cell-specific deletion of FcγRIIB leads to autoantibody production and a lupus phenotype (13, 14). FcγRIIB also plays an important role in aberrant responses to infection, such as to malaria (15).

Several studies have suggested a role for FcγRIIB in B cell tolerance, either through an effect on germinal center (GC) B cells, B-1 cells, or plasma cell (PC) survival (16-22). Mechanisms reported to contribute to the development of autoreactivity from the loss of FcγRIIB include activation of bystander autoreactive GC B cells, loss of follicular exclusion, and absence of PC apoptosis (23-25). Whereas most studies have focused on the role of FcγRIIB in the GC, recent developments have highlighted the role of extrafollicular B cell responses in SLE (26-28), necessitating an expanded analysis of the role of FcγRIIB on B cells. We analyzed the role of FcγRIIB in B cell-specific FcγRIIB-deficient mice, with a focus on extrafollicular responses. We show that loss of FcγRIIB in B cells leads to aberrant MZ activation and subsequent extrafollicular autoreactive PC responses.

## Results

### Increased spontaneous autoantibody IgG3 responses in Fcgr2b cKO mice

We first characterized the spontaneous autoantibody production in mice with a B-cell specific FcγRIIB condition knockout (*Fcgr2b* cKO). As reported previously, we observed increased anti-dsDNA IgG, which were present in mice around 4-5 months of age (Figure 1A, Figure S1A). Interestingly, when analysing the IgG subclasses, we observed a significant increase only in IgG3 anti-dsDNA (Figure 1B). Using a flow cytometry assay we developed to assess anti-nuclear antibody (ANA)-positive PCs (29), we observed an increased frequency of ANA+ IgG+ PCs in the spleen of *Fcgr2b* cKO mice (Figure 1C-D). In line with the serum data, the increase in ANA+ IgG+ PCs was only significant in the IgG3 subclass (Figure 1E). The frequency of ANA+ PCs within other isotypes or in the BM was not increased (Figure S1B-F).The increase in ANA+ IgG3+ PCs in the spleen suggested an extrafollicular B cell response.

**Figure 1:**
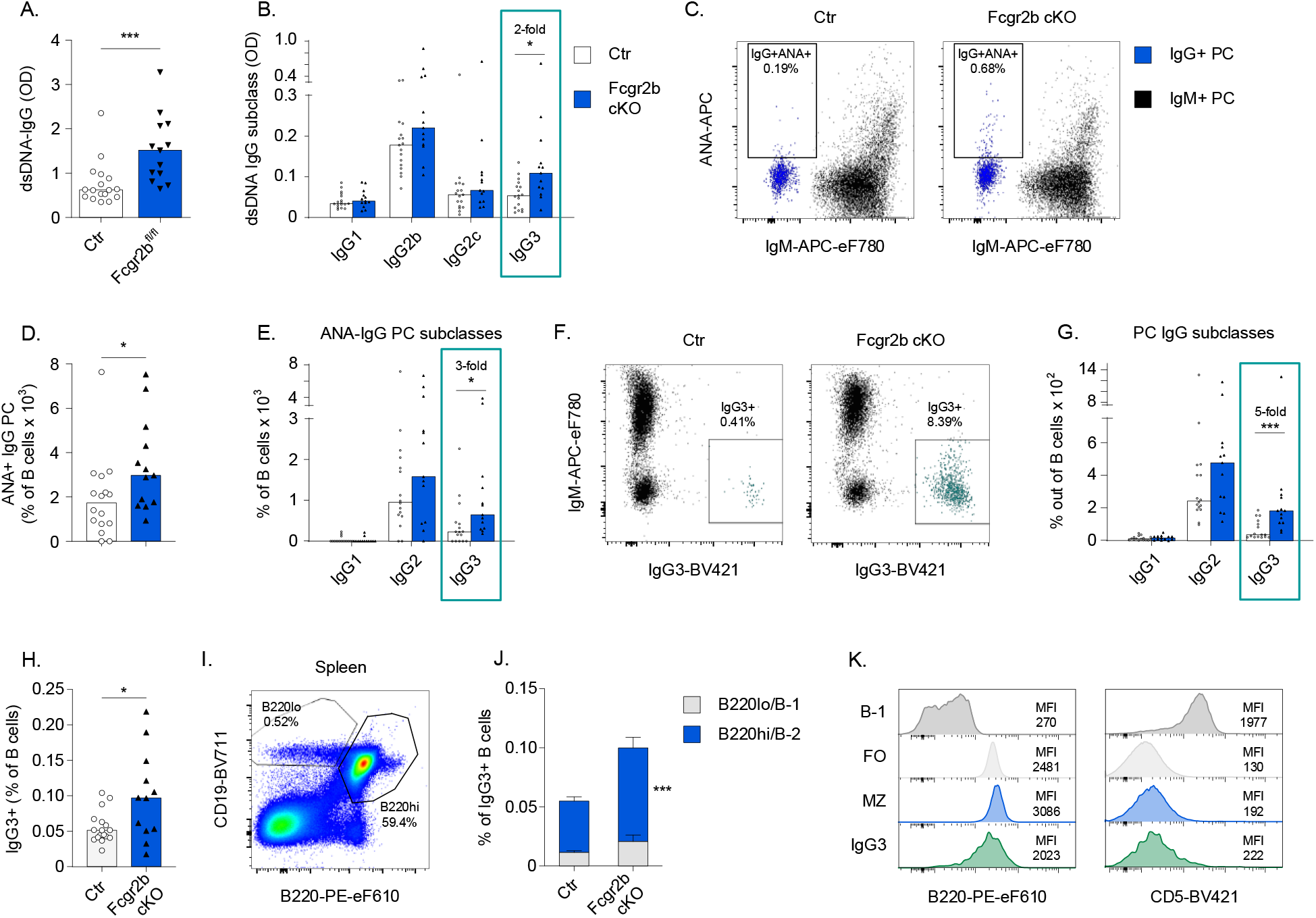
Increased spontaneous autoantibody IgG3 responses in *Fcgr2b* cKO mice Female control (Ctr) and *Fcgr2b* cKO mice were bred until 10-12 months of age after which (auto)antibodies in serum and PCs in spleen were characterized. ANA reactivity of PCs was established using flow cytometry. A,B) dsDNA ELISA for total IgG and IgG subclasses in serum of *Fcgr2b* cKO mice. C) Representative example of ANA staining in IgG and IgM PCs in spleen. D,E) Frequency of ANA+ IgG+ PCs in spleen; total IgG (D) and by IgG subclass (E). F) Representative example of IgM and IgG3 staining in total PCs. IgG3+ cells are indicated in green. G) Frequency of IgG+ PCs in spleen separated by subclass. H) Frequency of IgG3+ B cells in Ctr and *Fcgr2b* cKO mice. I) Representative example of staining strategy for B-1 and B-2 cells in spleen. J) Proportion of spleen IgG3+ B cells with a B-1 or B-2 phenotype respectively, gated as in H. K) Representative example of staining for B220 and CD5 in total IgG3+ B cells, compared to B-1, follicular (FO), and marginal zone (MZ) B cells in spleen. Data are shown as median with each symbol representing an individual mouse or mean +/-SEM (J) (n=12-17 per group pooled from 2-3 independent experiments). Asterisks indicate significant differences (*p<0.05; **p<0.01; ***p<0.001) obtained using Mann Whitney U test.

Since we previously showed that increased ANA+ IgG PCs in SLE patients and lupus-prone mice occur through aberrant IgG PC differentiation rather than an antigen-specific tolerance defect (29), we also analysed tolerance checkpoints for ANA+ B cells and PCs in *Fcgr2b* cKO mice. B cell-intrinsic FcγRIIB deficiency did not affect the percentage of ANA+ mature naïve B cells or more immature B cell subsets in BM or spleen (Figure S2A-C, Figure S3). In contrast, a specific increase in the frequency of spleen IgG+ PCs and serum levels of total IgG were found (Figure S1G-I, M, N). Again, the most prominent increase was in the IgG3 subclass, both in PCs and in serum titer (Figure 1F,G, Figure S1J-L, O-R). Together, these results indicate that spontaneous (auto-)antibody production occurs through enhanced differentiation or survival of IgG3 PCs.

*Fcgr2b* cKO spontaneously displayed an increased frequency of resting IgG3+ B cells (Figure 1H, Figure S4A-E). IgG3 is usually derived from B-1 cells or MZ B cells and increased numbers of peritoneal B-1 cells have been reported in complete *Fcgr2b*-/-mice (22). Most IgG3+ B cells in *Fcgr2b* cKO mice had a B-2 (CD19+B220hi) phenotype and were CD5-negative (Figure 1I-K). In addition, the frequencies of B-1a or B-1b cells in spleen and peritoneum were unaffected in *Fcgr2b* cKO mice (Figure S4F-M), whereas MZ B cell frequencies were increased in *Fcgr2b* cKO mice (Figure S4N,O).

### Increased extrafollicular PC responses

Since IgG3 has been associated primarily with PC responses derived from MZ or B-1 cells, we next analysed the extrafollicular response to immunization with prototypical T-independent and T-dependent antigens, NP-Ficoll and NP-CGG in alum, respectively. We observed a large increase in NP-IgG, but not NP-IgM, serum titers in *Fcgr2b* cKO mice, on day 7 following immunization with NP-Ficoll (Figure 2A). The anti-NP response of all IgG subclasses was significantly increased, with a marked increase in IgG3 (Figure 2B). In line with serum antibody levels, NP-specific IgG+ PCs in the spleen were increased, whereas no increase in IgM PCs was observed (Figure 2C-E). The greatest increase of NP-specific PCs was present in the IgG3 subclass (Figure 2F,G). Besides an increase in NP+ IgG3+ PCs, there was an increased frequency of NP+IgG3+ B cells in the spleen of *Fcgr2b* cKO mice (Figure 2H,I).

**Figure 2:**
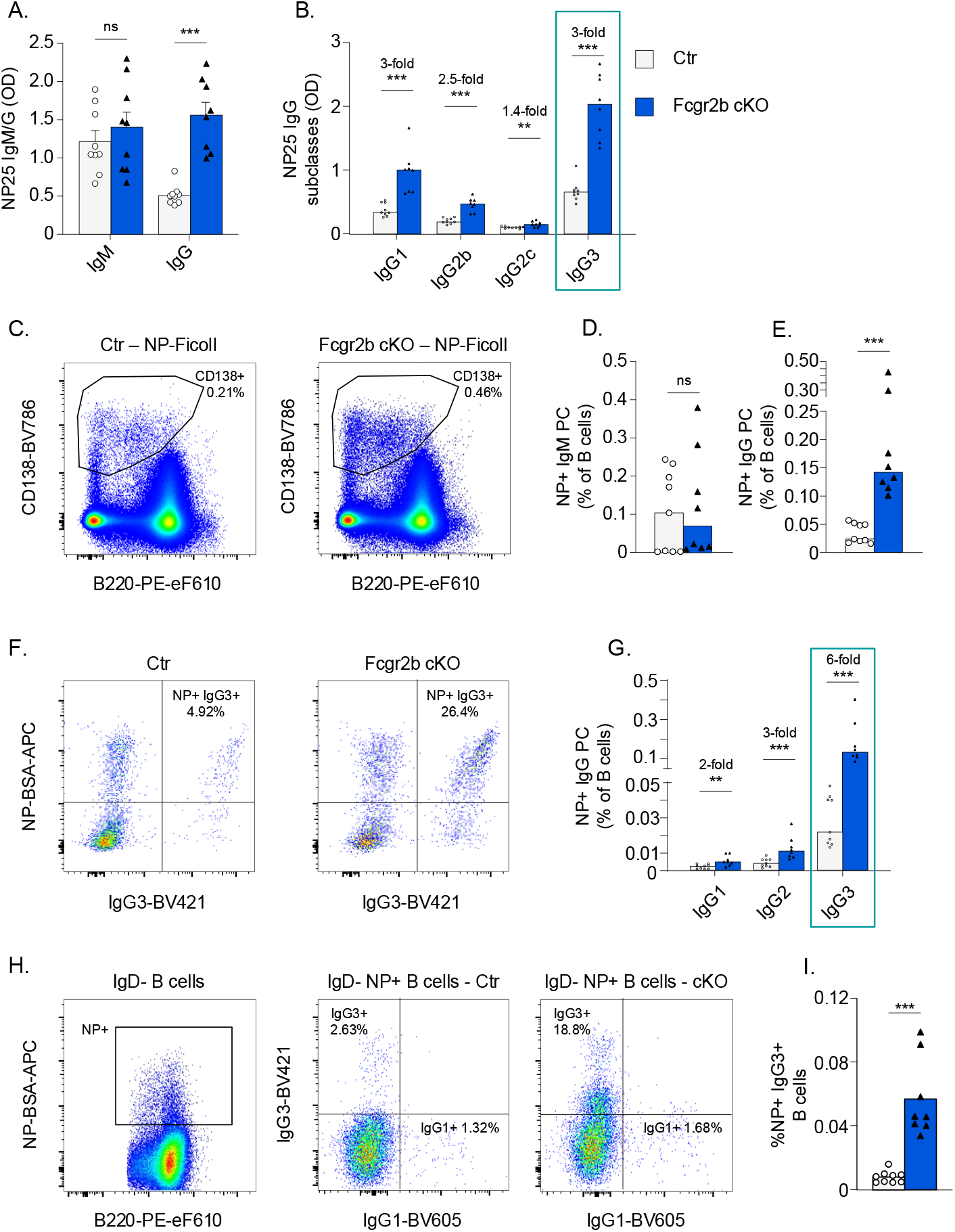
Increased extrafollicular responses following immunization. A-F) Female Ctr and *Fcgr2b* cKO mice were immunized with NP-Ficoll. Serum and splenocytes were obtained after 7 days. A,B) Levels of NP-specific antibodies, separated by isotype and subclass. C) Representative example of PC staining following NP-Ficoll immunization (concatenated data of 4×1k cells from 4 different mice in each group). D,E) Frequency of NP-specific PCs in spleen by isotype, as a percentage of B cells. F) Representative examples of intracellular IgG3 and NP staining in spleen PCs. G) Frequency of NP-specific PCs in spleen as percentage of B cells, separated by IgG subclass. H) Representative example of surface NP gating on IgD-B cells (left) and IgG1 and IgG3 staining in IgD-NP+ B cells (middle and right; ∼10k cells concatenated from 4 mice in each group). I) Frequency of IgG3+ NP+ B cells out of total B cells. Data are shown as median with each symbol representing an individual mouse (n=8-9 per group pooled from 2-3 independent experiments). Asterisks indicate significant differences (*p<0.05; **p<0.01; ***p<0.001) obtained using Mann whitney U test.

Whereas total NP-IgG and NP-IgM were not increased in *Fcgr2b* cKO mice following a T-dependent response to NP-CGG (Figure S5A), we observed an increase in NP-IgG3 serum levels and PCs (Figure S5B,C). Likewise, NP+ IgG3+ B cells were increased (Figure S5D,E). In line with an extrafollicular origin of NP-specific IgG3 following NP-CGG immunization, low-affinity NP-25 IgG3 levels peaked early (around day 14), and high-affinity NP-2 IgG3 levels were detected only at low levels at all time points up to day 42 (Figure S5F-G).

Together these data suggest that B cell specific FcγRIIB deficiency leads to greatly enhanced extrafollicular responses to immunization, with a particular strong increase in IgG3 responses that may be of MZ origin.

### Increased activation of MZ B cells in absence of FcγRIIB

Since MZ or B-1 cells could be the origin of increased extrafollicular responses in *Fcgr2b*CKO mice, expression of FcγRIIB was analysed in several B cell subsets in mice (Figure 3A,B). Expression of FcγRIIB was highest in MZ B cells and IgG3+ B cells, compared to B-1 cells and follicular (FO) B cells. Following immunization with NP-Ficoll and NP-chicken gamma globulin (NP-CGG), the highest expression of FcγRIIB was observed in NP+ IgG3+ B cells, compared to naïve B cells or IgG1+ NP+ B cells (Figure S6A-C). These results suggest that MZ B cells and IgG3+ B cells may be most susceptible to inhibition by FcγRIIB; stated otherwise: MZ B cells may be most activated in *Fcgr2b* cKO mice.

**Figure 3:**
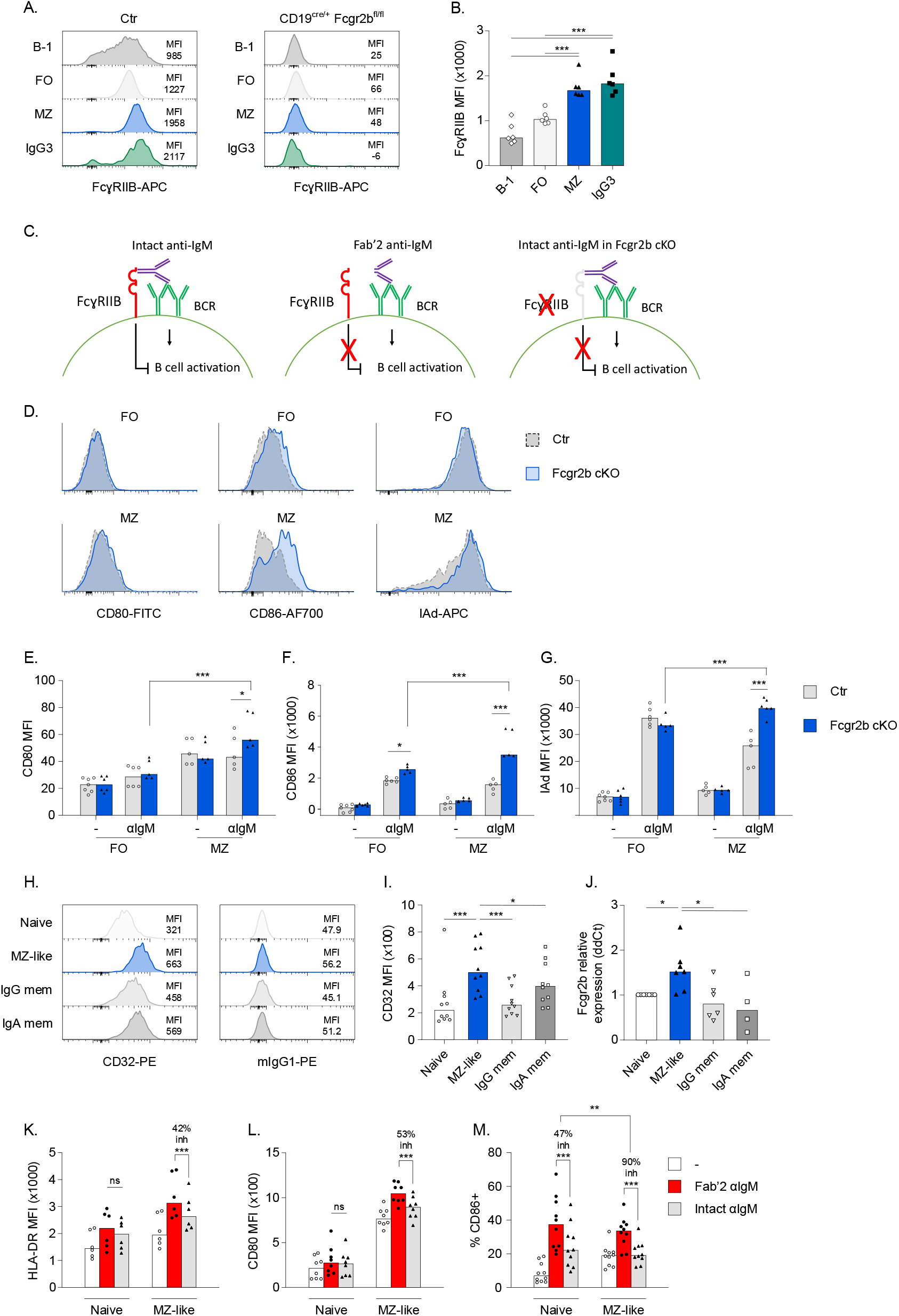
Phenotype and activation of MZ B cells in *Fcgr2b* cKO mice and humans. A) Representative example of staining for FcγRIIB in different B cell subsets and IgG3+ B cells in the spleen. Panel on the right shows staining in *Fcgr2b* cKO cells. Similar results were obtained using isotype controls (data not shown). B) FcγRIIB staining intensity in different B cell subsets from control mice. C) Schematic of experimental approach to investigate the effect of FcγRIIB on B cell activation. Using intact anti-IgM, both BCR and FcγRIIB are engaged (left); using Fab’2 anti-IgM (middle) or *Fcgr2b* cKO cells (right), only BCR is engaged. D-G) FO and MZ B cells were sorted from the spleens of Ctr and *Fcgr2b* cKO mice, followed by stimulation with 3 ug/mL anti-IgM for 20 hours. Activation was measured by flow cytometry. Representative examples (D) and summary (E-G) of expression of CD80, CD86 and IAd. H,I) Representative examples and summary of expression of CD32 analysed by flow cytometry in different human B cell subsets, gated as shown in Figure S8A. J) qPCR for *FCGR2B* in sorted human B cell subsets. Relative expression was normalized to polr2a, after which ddCt was calculated compared to Naïve B cells. K-M) Sorted human naïve and MZ-like B cells were stimulated with 3 ug/mL intact anti-IgM or equimolar concentrations (2 ug/mL) of Fab’2 anti-IgM 20 hours or were left untreated as controls. Upregulation of HLA-DR, CD80 and CD86 was measured by flow cytometry. Percentage inhibition was calculated for each donor, (the median % inhibition is indicated in the figure). Data are shown as median with each symbol representing an individual (mouse or human) (n=6 per group for B; n=5-7 per group for D-F; n=6-10 per group for I-M; each pooled from 2-5 independent experiments; except B which was from 1 experiment). Asterisks indicate significant differences (*p<0.05; **p<0.01; ***p<0.001) obtained using One-way ANOVA with Bonferroni posthoc test (B, I,J), or Two-way ANOVA with Bonferroni posthoc test (E-G, K-M).

Since we observed increased expression of FcγRIIB in MZ B cells, we asked if there was a differential inhibitory effect of FcγRIIB on FO B cells and MZ B cells. Sorted B cell subsets were activated through BCR crosslinking in the presence (intact IgM crosslinking antibodies) or absence of FcγRIIB engagement (*Fcgr2b* cKO cells or Fab’2 IgM crosslinking antibodies) (Figure 3C). We first determined optimal concentrations by Fab’2 IgM crosslinking antibodies for upregulation of CD80, CD86 and MHC class II (IAd) in FO B cells (Figure S6D-F), and observed increased upregulation of CD80 and CD86 in MZ B cells compared to FO B cells (Figure S6G-I). We then analysed the effect of FcγRIIB engagement on BCR-mediated activation of FO and MZ B cells. FcγRIIB deficiency increased activation of MZ B cells, whereas it had almost no effect in FO B cells (Figure 3D-G). The absence of a strong effect on FO B cells was surprising, so we confirmed the observation by comparing equimolar concentrations of intact versus Fab’2 anti-IgM antibodies in control mice (Figure S6J-L).

To address these findings in humans, we analyzed expression and inhibitory function of FcγRIIB in various B cell subsets from healthy donors. IgM+ CD27+ B cells have been proposed as the human equivalent of MZ B cells in mice (MZ-like B cells) (30). We isolated these cells by FACS and confirmed their MZ-like phenotype by high *SOX7* and low *HOPX* expression, relative to conventional naive and IgG+ memory B cells (30) (Figure S7A-B). Next, we analyzed expression of FcγRIIB in MZ-like B cells, naïve B cells, and IgG and IgA memory B cells. As in mice, the human MZ-like compartment had the highest expression of FcγRIIB, on both the protein and RNA level (Figure 3H-J). MZ-like B cells were also more strongly activated than naïve B cells by BCR crosslinking using Fab’2 anti-IgM across a range of concentrations (Figure S7C-E). We next analysed the effect of FcγRIIB-BCR crosslinking in MZ-like B cells compared to naïve B cells, and observed a stronger inhibitory effect of FcγRIIB engagement in MZ-like B cells than on naive B cells (Figure 3K-M, Figure S7I). (For example, while CD86 upregulation by BCR engagement was similar between naïve B cells and MZ B cells (Figure S7H), the inhibitory effect of FcγRIIB in in MZ-like B cells returned expression close to baseline levels (Figure 3M)). Upregulation of HLA-DR and CD80 upon BCR crosslinking was also stronger in MZ-like B cells (Figure S7F,G), and FcγRIIB-mediated inhibition was most pronounced in this subset (Figure 3K-L).

MZ-like B cells have been reported to be more prone to activation; we now show that they also exhibit stronger inhibition of activation through FcγRIIB, an effect that is present in both mice and humans.

### FcγRIIB inhibits MZ B cell activation through inhibition of Erk phosphorylation and calcium flux

We next analysed phosphorylation of signaling molecules downstream of the BCR and FcγRIIB (Figure 4A). We chose to study Syk, which is upstream of FcγRIIB signaling, and therefore should not be affected, as well as Erk1/2 which is downstream of FcγRIIB signaling and is important for B cell activation and plasma cell differentiation (31). Consistent with the increased upregulation of activation markers, we found increased phosphorylation of Syk and Erk1/2 in murine MZ B cells compared to FO B cells stimulated with Fab’2 anti-IgM (Figure 4C-D). There was strong inhibition of Erk1/2 phosphorylation in MZ B cells in the presence of *Fcgr2b* engagement with intact anti-IgM antibodies, but no inhibition of Syk phosphorylation (Figure 4B-D). A significant decrease in Erk1/2 phosphorylation was only observed in MZ B cells and not in FO B cells (Figure 4D). Consistent with this observation, Erk1/2 phosphorylation was increased in the absence of FcgRIIb engagement in *Fcgr2b* cKO mice compared to control mice (Figure 4E,F). Again, this difference was only observed in MZ B cells.

**Figure 4:**
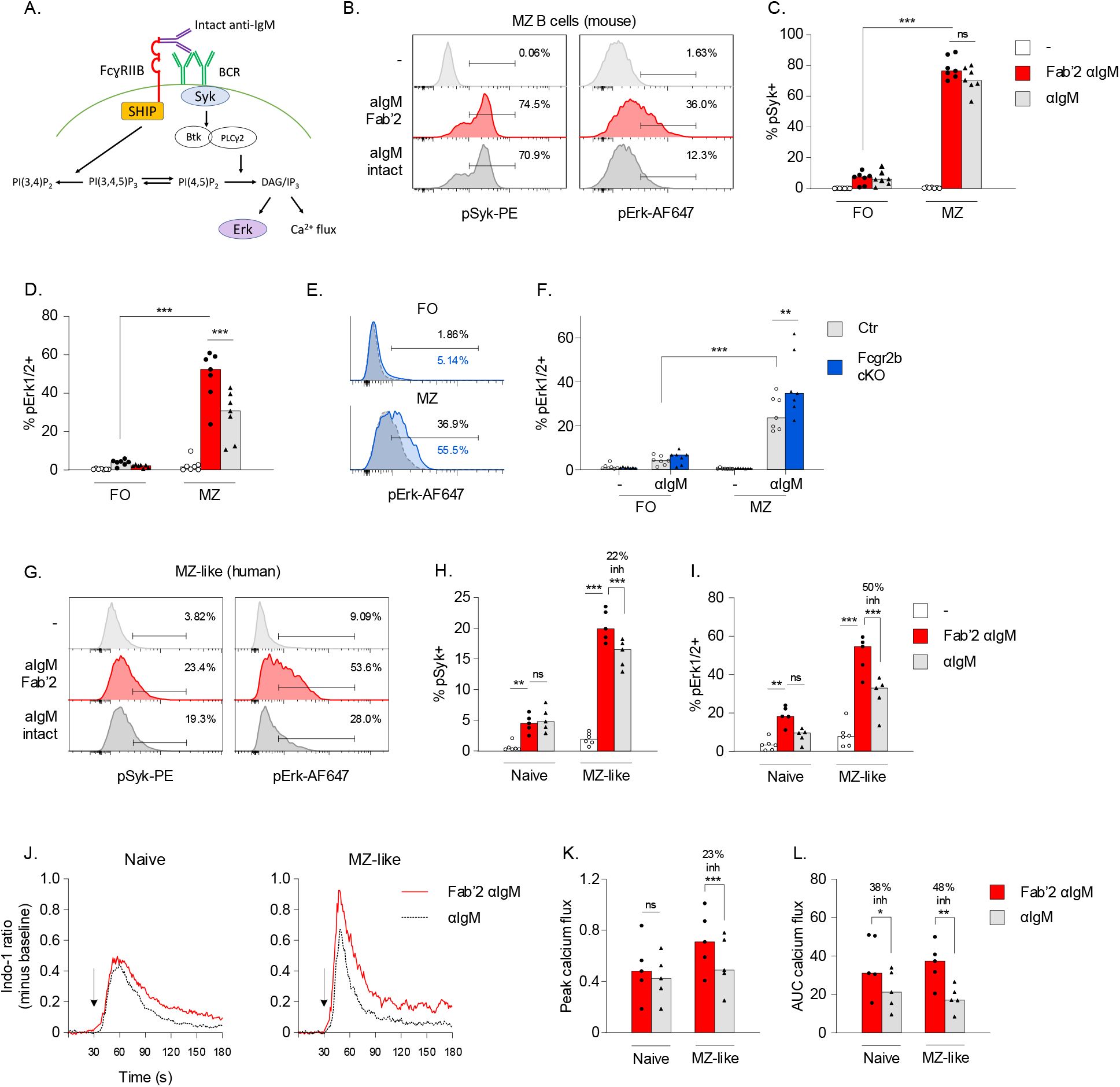
The effect of FcγRIIB on B cell signaling in MZ B cells. A) Simplified schematic diagram of key signaling molecules downstream of the BCR and FcγRIIB. B-D) Representative examples and summary of FO and MZ B cells from Ctr mice which were stimulated with 3 ug/mL intact anti-IgM or equimolar concentrations (2 ug/mL) of Fab’2 anti-IgM for 10 min, after which the phosphorylation of signaling molecules was analysed by phosphoflow. E,F) Representative examples and summary of pErk expression in FO and MZ B cells from Ctr and *Fcgr2b* cKO mice which were stimulated with 3 ug/mL intact anti-IgM as described in B-D. G-I) PBMCs from healthy donors were stimulated with 3 ug/mL intact anti-IgM or equimolar concentrations (2 ug/mL) of Fab’2 anti-IgM for 60 min, after which the phosphorylation of signaling molecules was analysed by phosphoflow. G) Representative example of Syk and Erk phosphorylation in MZ-like B cells gated as in Figure S7A. H,I) Comparison of intact versus Fab’2 anti-IgM in naïve and MZ-like B cells. Median % inhibition calculated per donor is indicated in the Figure. J-L) PBMCs from healthy donors were labeled with Indo-1 for calcium flux measurements. After 30 seconds of baseline measurement, 75 ug/mL intact anti-IgM or equimolar concentrations (50 ug/mL) of Fab’2 anti-IgM were added and the measurement continued for 2.5 min. Peak calcium flux and area under the curve (AUC) were calculated using Flowjo. J) Representative examples of calcium flux in naïve and MZ-like B cells. K,L) Comparison of peak and AUC of calcium flux following intact versus Fab’2 anti-IgM in naïve and MZ-like B cells. Data are shown as median with each symbol representing an individual mouse (n=4-7 per group for B-D; n=5-6 per group for G-L; each pooled from 2-3 independent experiments). Asterisks indicate significant differences (*p<0.05; **p<0.01; ***p<0.001) obtained using Two-way ANOVA with Bonferroni posthoc test.

In human MZ-like B cells, we similarly observed increased phosphorylation of Syk and Erk1/2 following BCR triggering in MZ-like cells compared to naive B cells (Figure S8A-B). Whereas a small inhibition of Syk phosphorylation was observed, FcγRIIB engagement strongly inhibited pErk in MZ-like B cells, but not naïve B cells (Fig. 4G-I). MZ-like B cells compared to naive B cells exhibited increased calcium flux upon BCR triggering (Figure S9), which was more strongly inhibited by FcγRIIB in MZ-like B cells than naive B cells, in particular the peak flux (Figure 4J-L).

These results point to a strong inhibitory effect of FcγRIIB on activation of MZ B cells through decreased phosphorylation of Erk1/2 and decreased calcium flux.

### IgG3 responses in Fcgr2b cKO mice are derived from MZ B cells

Because our results suggest that MZ B cells are a likely source for the increased IgG3 response in *Fcgr2b* cKO mice we generated double KO (dKO) mice that lack Notch2 and FcγRIIB in B cells, resulting in large reductions in MZ B cells (32) and B cell-specific FcγRIIB deficiency. These mice exhibited deficiency of MZ B cells (∼80% reduction), whereas FO B cells and B-1 cell numbers were maintained (Figure 5A,B, Figure S9A-C). Similar to what has been described (33), *Notch2* cKO mice exhibited decreased NP-specific IgM and IgG serum levels and PCs, but there was no effect of FcγRIIB on the IgM response, both in presence or absence of Notch2 (Figure S9D,F). More importantly, the increased IgG response to NP-Ficoll that was observed in *Fcgr2b* cKO was completely reversed in dKO mice lacking both Notch2 and FcγRIIB in B cells (Figure S9E,G). Furthermore, the increase in NP-IgG3 serum titers and NP+ IgG3+ B cells and PCs generated by FcγRIIB deficiency was absent in dKO mice (Figure 5C-F). IgG3+ B-2 B cells, but not IgG3+ B-1 B cells, were significantly reduced in MZ-deficient dKO mice (Figure S9H,I). Together, these results show that MZ B cells are responsible for enhanced extrafollicular antibody responses in *Fcgr2b* cKO mice.

**Figure 5:**
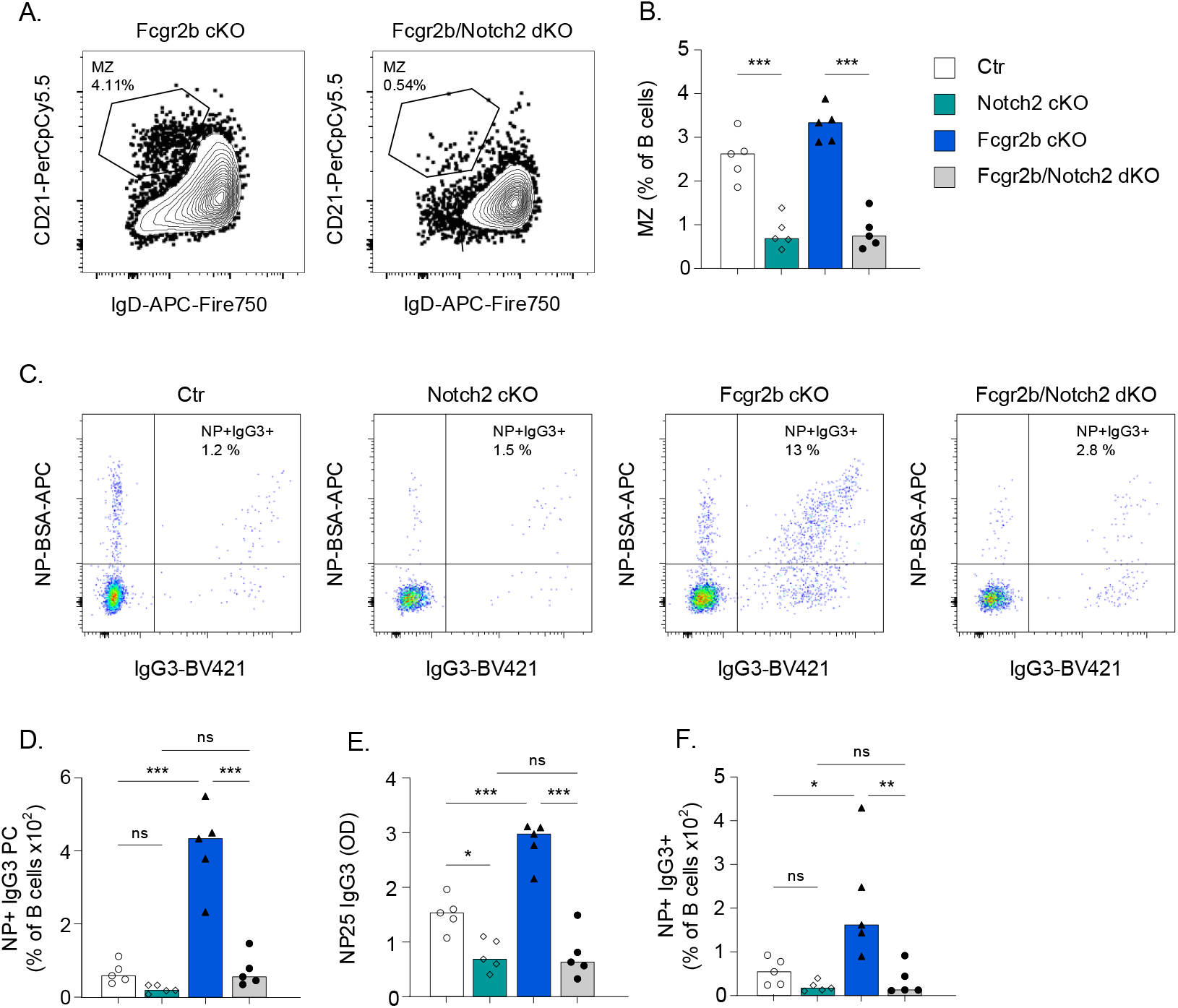
Combined FcγRIIB and MZ deficiency reverses enhanced extrafollicular responses. Female Ctr, *Notch2* cKO, *Fcgr2b* cKO, and *Notch2*/*Fcgr2b* dKO mice were immunized with NP-Ficoll. Serum and splenocytes were obtained after 7 days. A,B) Representative examples of MZ B cell frequencies in *Fcgr2b* cKO and *Notch2*/*Fcgr2b* dKO mice. C) Representative examples of intracellular IgG3 and NP staining in spleen PCs. D) Frequency of NP-specific IgG3+ PCs in spleen, as a percentage of B cells. E) Levels of NP-specific IgG3 in serum. F) Frequency of NP-specific IgG3+ B cells in spleen. Data are shown as median with each symbol representing an individual mouse (n=5 per group). Asterisks indicate significant differences (*p<0.05; **p<0.01; ***p<0.001) obtained using Two-way ANOVA with Bonferroni posthoc test.

In summary, our results show increased extrafollicular responses in B cell-intrinsic FcγRIIB-deficient mice characterized by a major increase in MZ-derived IgG3 responses. MZ B cells are highly sensitive to inhibition through FcγRIIB, which can be explained by the high expression of FcγRIIB in MZ B cells compared to other B cell subsets. High FcγRIIB expression results in strong inhibitory signaling through FcγRIIB in MZ B cells, an effect that was observed in both mice and humans.

## Discussion

In this study, we analyzed B cell tolerance and extrafollicular PC differentiation in B cell intrinsic FcγRIIB-deficient mice (14). We have identified spontaneous PC differentiation in *Fcgr2b* cKO mice, leading to increased serum autoantibody levels, in particular of the IgG3 isotype. In addition, *Fcgr2b* cKO mice showed increased IgG3 responses following immunization, which was MZ B cell-dependent. In both mice and humans, MZ B cells showed the highest expression of FcγRIIB; this high expression associates with increased FcγRIIB-mediated inhibitory effects on Erk phosphorylation and calcium signaling in vitro. Thus, B cell-intrinsic FcγRIIB-deficiency is linked to increased extrafollicular responses through an effect specifically on MZ B cells.

Similar to previous studies on lupus prone mice performed in our laboratory (29), we found that B cell deficiency of FcγRIIB led to an increase in PCs, without an increase in the fraction of ANA+ IgG PCs, suggestive of increased B cell activation and PC differentiation rather than an antigen specific tolerance defect. This is also in line with the increased responses we observed to immunization with NP-Ficoll and NP-CGG in *Fcgr2b* cKO mice. Of note, these mice did not develop a fulminant lupus phenotype despite the presence of autoreactive PCs in the spleen and autoantibodies in serum. Complete *Fcgr2b* KO mice on the C57BL/6 background spontaneously develop autoantibody titers and develop a fulminant lupus phenotype (9, 14), probably due to combined effects on B cells and myeloid cells (34).

In contrast to studies with the full FcγRIIB KO (22), we did not observe an effect of B cell-intrinsic FcγRIIB-deficiency on B-1 numbers in spleen or peritoneum, and FcγRIIB expression in B-1 cells was low, suggesting that the effect on the B-1 compartment is indirect. We now show that the enhancement of NP-Ficoll responses in *Fcgr2b* cKO mice is completely reversed in *Notch2*/*Fcgr2b* deficient dKO mice. Since these mice had largely reduced MZ numbers without significant reductions in B-1 cells (32), this suggests that enhanced extrafollicular responses to NP-Ficoll are dependent on MZ B cells.

As has been reported in the literature, we observed increased BCR-mediated activation of MZ B cells compared to FO or naïve B cells, in both humans and mice (35). Importantly, we showed that MZ B cells also had stronger inhibition mediated by FcγRIIB co-engagement. In the MZ B cell compartment, FcRIIB-mediated inhibition was linked to diminished Erk phosphorylation and calcium flux. Previous studies into the effect of FcγRIIB-BCR crosslinking on B cell signaling have revealed quite variable effects, probably depending on the source and B cell subset studied. Most studies have shown no effect on Syk phosphorylation (36, 37). FcγRIIB engagement preferentially recruits SHIP, which acts to hydrolyze phosphatidylinositol (3,4,5)P3 and inositol(1,3,4,5)P4, downstream of Syk (38). *Fcgr2b* engagement can also inhibit B cell responses in SHIP-independent manners (such as through SHP-1/SHP-2 recruitment), maybe explaining the weak inhibition of Syk that we observed in the presence of FcγRIIB engagement in human MZ-like cells (39). FcγRIIB-BCR crosslinking also leads to earlier closing of calcium channels (37, 40), consistent with our data showing diminished calcium flux. Since we were interested in the difference between MZ B cells and FO B cells, we used primary B cells as opposed to cell lines which were used in most previous studies. Our results show the importance of analyzing signaling in a cell type specific manner, as different B cell subsets can respond quite differently, and with our studies we were able to reveal strong inhibitory signaling of FcγRIIB in MZ B cells.

*FCGR2B* risk alleles which lead to diminished expression and/or inhibitory function predispose to SLE. Furthermore, diminished expression of FcγRIIB on memory B cells and plasmablasts from SLE patients has been shown irrespective of the presence of risk alleles (41, 42). We and others have recently demonstrated that most SLE patients exhibit enhanced PC differentiation and some exhibit increased extrafollicular PC differentiation (26, 28, 43). Our current data show how diminished expression of FcγRIIB can lead to increased extrafollicular B cell activation and autoantibody production. Since MZ B cells are known to have increased autoreactivity (29, 44-46), a loss in the regulation of extrafollicular MZ B cell responses through diminished function of FcγRIIB may therefore lead to autoantibody production. In summary, we present a model where high expression of FcγRIIB in MZ B cells is necessary to prevent ongoing activation, and thereby functions as a feedback loop to prevent ensuing autoimmunity, in particular through its effect on extrafollicular MZ responses. These data also suggest that SLE patients with a risk allele for *FCGR2B* may have a specific impairment in MZ B cells, and may be those with most fluctuation in autoantibody titers. Understanding how risk alleles predispose to autoantibody production has implications for precision medicine.

## Materials and methods

### Mice

*Fcgr2b*^flox/flox^ mice were a kind gift from Jeffrey Ravetch (14). CD19^cre^ mice (stock number 006785), B6.C-H2d (stock number 000359) were obtained from Jackson and were crossed with *Fcgr2b*^fl/fl^ mice to generate CD19^cre/+^ *Fcgr2b*^fl/fl^ H2d/d (*Fcgr2b* cKO) mice. Control mice were either carrying only the Cre allele (CD19^cre/+^ *Fcgr2b*^wt/wt^ H2d/d), only the flox alleles (CD19^+/+^ *Fcgr2b*^fl/fl^ H2d/d). To generate MZ-deficient mice, CD19^cre/+^ control mice and *Fcgr2b* cKO mice were crossed with *Notch2*^flox/flox^ (Jackson stock number 010525) to generate CD19^cre/+^ *Notch2*^fl/fl^ H2d/d and CD19^cre/+^ *Fcgr2b*^fl/fl^ *Notch2*^flox/flox^ H2d/d mice.

For analysis of spontaneous autoimmunity, mice were kept until 10-12 months of age. For analysis of immunization responses, 8-16 week old female mice were immunized i.p. with 50 ug NP-Ficoll (conjugation ratio 55:1) in 100 uL saline (Biosearch Technologies) or 100 ug NP-CGG (conjugation ratio 20-29:1) (Biosearch Technologies) in Imject Alum (Thermofisher), and followed for 7 days (both NP-Ficoll and NP-CGG) and for 42 days (NP-CGG only). For functional in vitro studies of FcγRIIB, spleens of 8-16 week old female mice were used.

Spleens and bone marrow was collected at the indicated timepoints followed by formation of a single cell suspension by mashing over a 70 um cell strainer. Red blood cell (RBC) lysis was performed using RBC lysis buffer (Biolegend). Peritoneum cells were obtained by injecting 10 mL HBSS + 5% FBS, followed by massaging the peritoneum, and withdrawal of the buffer with cells. Serum was obtained through submandibular bleeding followed by centrifugation.

### Human subjects

Buffy coats from healthy donors were obtained through Sanquin blood bank (The Netherlands). PBMCs were isolated using standard Ficoll procedure. For signaling experiments, frozen PBMCs obtained from buffy coats were used.

### ELISA

Half-area ELISA Plates (Corning) were coated with 10 ug/mL of NP2-, or NP25-BSA (Biosearch technologies), anti-mouse IgG, IgM, IgG1, IgG2b, IgG2c, or IgG3 unlabeled antibody (Southern Biotech) overnight at 4°C, or with 100-400 ug/mL sonicated filtered calf-thymus DNA (Calbiochem) overnight at 37 degrees C uncovered. Plates were washed with PBS .05% Tween 20 and blocked with 1% BSA in PBS. Diluted serum samples were incubated for 1.5 hours. Serum samples for anti-DNA IgG and IgG subclasses were diluted 1:100. Serum samples from NP-immunized mice were diluted 1:10.000 (day 7) or 1:50.000 (timecourse day 0, 14, 28, 42) for NP-specific IgM and IgG, and 1:2.500 for NP-specific IgG subclasses. Plates were washed with wash buffer and secondary goat polyclonal AP-labeled anti-mouse IgG, IgM, IgG1, IgG2b, IgG2c, or IgG3 (Southern Biotech) was added for 1 hour. After washing, plates were developed using phosphatase substrate (Sigma) dissolved in distilled water with 50 mM NaHCO3 (Sigma) and 1 mM MgCl2 (Sigma). Plates were read at a wavelength of 405 nM on a 1430 Multilabel Counter Spectrometer (PerkinElmer).

### Flow Cytometry

For flow cytometry phenotyping, cells were pre-incubated on ice for 5 min with Fc block (anti-mouse or anti-human) for each staining, except staining of FcγRIIB. After this, cells were stained for 30 minutes with antibodies to cell membrane proteins followed by washing in 5% Fetal Bovine Serum (FBS) in Hank’s Balanced Salt Solution (HBSS) or 0.5% BSA in PBS. For surface staining, cells were washed and stored in 1% PFA until acquisition. For intracellular staining, cells were fixed and permeabilized with Foxp3 transcription factor fixation/permeabilization kit (eBioscience) for 45 minutes on ice. After fixation and permeabilization, the intracellular staining cocktail was incubated for 30 min in Permeabilization buffer to visualize immunoglobulins and ANA or nitrophenol (NP) reactivity in PCs.

Antibodies used are listed in Table S1. For staining of NP-specific B cells in immunized mice, NP(6)-BSA-biotin (Biosearch Technologies) was pre-incubated for 30 min with streptavidin-APC in a 1:1 molar ratio, after which cells were incubated with 0.1 µg/mL NP-BSA-biotin + streptavidin-APC during either the surface or intracellular staining step. Surface and intracellular ANA staining was performed as described (29).

Mouse gating strategies were based on our prior studies (26, 29). Live cells were gated using DAPI or fixable viability dye eF506. B cell subsets were as follows in mouse bone marrow: Immature B cells: B220+ CD43-CD24hi CD21lo IgDlo IgMhi; Emigrating B cells: B220+ CD43-CD24hi CD21lo IgDhi IgMhi; Recirculating B cells: B220+ CD43-CD24+ CD21+ IgMhi IgDhi. B cell subsets in mouse spleen were: T1: B220hi CD93+ CD23lo IgMhi; T2: B220hi CD93+ CD23hi IgMhi; T3: B220hi CD93+ CD23hi IgMlo; FO: B220hi CD93-CD21lo with either IgDhi, CD23+ or both; MZ: B220hi CD93-CD21high with either IgDlo, CD23lo or both. B1 B cells in spleen and peritoneum were gated as: B220lo CD19hi CD93-CD21-IgDlo, with CD5 used to separate B1a and B1b cells. Memory B cells were gated as IgD-CD38+ GL7, Activated/pre-GC B cells were gated as IgD-CD38+ GL7+, and GC B cells were gated as IgD-CD38-GL7hi. CD95 expression on GC B cells was used to further confirm the GC gating. PCs were gated based on B220lo/+ and CD138high and were validated by intracellular immunoglobulin staining. IgG subclasses and specificity (ANA/NP) were determined within memory, activated, GC B cells and PCs populations. For Phosflow experiments, staining for CD93, CD21 and CD23 was performed prior to activation (due to loss of signal following fixation/permeabilization), while all other antibodies were included in the intracellular staining mix.

For human B cell gating strategies all cells were gated as Live based on fixable viability dye eF506. Naïve B cells were gated as CD19+ CD27-CD38lo IgG-IgA-IgDhi IgMlo; MZ-like B cells were gated as CD19+ CD27+ CD38lo IgG-IgA-IgDlo IgMhi. For functional assays, staining with IgM/IgD needed to be avoided to prevent activation of cells through their BCR, but their phenotype was confirmed by subsequent staining for IgM/IgD. For Phosflow experiments, CD27 staining was performed prior to activation (due to loss of signal following fixation/permeabilization), while all other antibodies were included in the intracellular staining mix.

### In vitro B cell Activation

For mouse in vitro activation studies, FO B cells and MZ B cells were sorted using the FACS Aria or Aria Sorp (BD). For dead cell exclusion, 20 ng/mL DAPI (Thermo) was added to the cells just prior to sorting. FO B cells and MZ B cells were cultured in RPMI medium + Pen/Strep + L-glutamine + 10% FBS and stimulated with polyclonal goat anti-mouse IgM (Southern Biotech; 0.6-150 ug/mL), or equimolar concentrations of the Fab’2 fragments (Southern Biotech; 0.4-100 ug/mL) for 20 hours after which activation was measured using flow cytometry as described above.

For human in vitro activation studies, naive and MZ-like B cells were sorted using BD FACS Aria. Cells were cultured in RPMI medium + Pen/Strep + L-glutamine + 10% FBS and stimulated with polyclonal goat anti-human IgM (Jackson Immunoresearch Affinipure; 0.6-75 ug/mL), or equimolar concentrations of the Fab’2 fragments (Jackson Immunoresearch Affinipure; 0.4-50 ug/mL) for 20 hours after which activation was measured using flow cytometry as described above.

### B cell Signaling

For Phosflow experiments, mouse splenocytes or human PBMCs were stimulated for 2, 10, and 60 min in RPMI medium + Pen/Strep + L-glutamine + 10% FBS, with the anti-IgM antibodies described above (intact 3-15 ug/mL; Fab’2 2-10 ug/mL). Cells were immediately fixed in 1x Phosflow Lyse/Fix buffer (BD). After 10-12 min incubation at 37 degr, cells were washed and permeabilized using Phosflow Perm Wash I (BD) for 30-60 min. Following a 5 min pre-incubation with Fc block (BD), cells were then incubated with a mix of antibodies to cell surface and phospho-specific signaling molecules or isotype controls at room temperature for 60 min.

For calcium flux experiments, human PBMCs were labeled with 0.5 uM Indo-1 AM (Thermo) in the presence of 0.02% Pluronic F-127 (Thermo) in IMDM medium + 2% FBS, for 45 min at 37 degr C. After that, cells were washed twice, followed by staining of surface markers on ice. After washing, cells were taken up in colorless IMDM + 2% FBS. Cells and stimuli were pre-warmed at 37 degrees for a minimum of 10 min before measuring. Calcium flux measurements were performed on LSR-II (BD). After 30 seconds of establishment of baseline Indo-1 signals, cells were stimulated with the anti-IgM antibodies described above (intact 15-75 ug/mL; Fab’2 10-50 ug/mL), after which acquisition was immediately resumed, and continued for 2.5 min. Ionomycin (Sigma) was used at 1 ug/mL as positive control.

### RNA isolation and qPCR

Human naïve, MZ-like, and conventional memory (IgG+ and IgA+) B cells were sorted according to the gating strategy described above. 50-200k cells per population were lysed in RLT buffer (Qiagen) with 10 uL/mL 2-ME (Merck) after which RNA was isolated using RNeasy Microprep. cDNA was synthesized using iScript (Biorad). qPCR was performed after pre-amplification using TaqMan PreAmp Master Mix (Thermo) and the Taqman assays mentioned below. qPCR was performed with TaqMan Fast Advanced Master Mix (Thermo) and multiplexed VIC and FAM Taqman assays (all from Applied Biosystems/Thermo): *POLR2A* VIC-MGB (Hs00172187_m1), *ACTB* VIC-MGB (Hs01060665_g1), *HOPX* FAM-MGB (Hs04188695_m1), *SOX7* FAM-MGB (Hs00846731_s1), *FCGR2B* FAM-MGB (Hs00269610_m1). qPCR was run on a CFX Opus machine (Biorad) using recommended cycling conditions.

### Data analysis

Analysis of flow cytometry data was performed using BD FACS DIVA and Flowjo. Graphing of calcium flux was performed using Graphpad after export of the binned data of each population in Flowjo. Cells were gated as % positive if there were two clear populations, whereas MFI was used when an entire population showed a shift. All MFIs shown are median fluorescence intensities. To control for differences in autofluorescent background or non-specific binding, isotype controls were used for in vitro activation and phosflow experiments. No difference in isotype control background were found between the populations that were compared. In vitro activation and phosphorylation data are shown for 3 ug/mL intact and 2 ug/mL Fab’2 anti-IgM antibodies. For both mouse and human studies, similar results were obtained with 10 and 15 ug/mL respectively (data not shown). For in vitro activation and signaling, % inhibition with intact anti-IgM was calculated per donor as: intact anti-IgM/(Fab’2 anti-IgM – unstimulated) * 100. Immunized mice were excluded when they exhibited no serum antibody response compared to baseline (n=1 for timeline NP-CGG response; n=3 for day 7 NP-CGG response; n=1 for day 7 NP-Ficoll response). One sample for FcγRIIB expression in human B cells was excluded from qPCR analysis because it was an extreme outlier in protein expression in naïve B cells (Figure 3I).

Statistical analysis was performed using Graphpad. For comparison of ex vivo mouse data with two groups, Mann Whitney Test was performed. For comparison of multiple groups, One-Way ANOVA with Bonferroni posthoc test was used. For analysis of in vitro activation with two groups, a paired T-test was used, and for comparison of two categorical variables, Two-Way ANOVA with Bonferroni posthoc test was used. P values <0.05 were considered statistically significant.

### Study approval

Mice were housed in according to AAALAC regulations and all mouse studies were approved the Institutional Animal Care and Use Committee (The Feinstein Institutes for Medical Research/Northwell health).

Studies with buffy coats from healthy donors were performed in accordance with the declaration of Helsinki and written informed consent was obtained from all donors.

## Author contributions

ANB designed and conducted experiments, analyzed data, and wrote the initial manuscript. SM designed and conducted experiments, and analyzed data and revised the manuscript. ALD conducted experiments and analyzed data. JS conceptualized the study, designed and conducted experiments, analyzed data, and wrote the manuscript. BD conceptualized the study, designed experiments, interpreted the data and wrote the manuscript. The order of the co–first authors was determined by who initiated the study (ANB) and who completed the study (SM).

## Acknowledgements

The authors thank the flow cytometry core facilities of the Feinstein Institutes for Medical Research and Leiden University Medical Center for their assistance in FACS sorting experiments. JS received financial support from American Autoimmune Related Disease Association. This work was further supported by NIH 1P01 AI073693.

## Notes

**Conflict of interest statement** The authors have declared that no conflict of interest exists.

### Competing Interest Statement

The authors have declared no competing interest.

