## Supplemental data for "FcγRIIB regulates (auto)antibody responses by limiting marginal zone B cell activation"

### Supplementary file (Figure S1-9, Table S1)

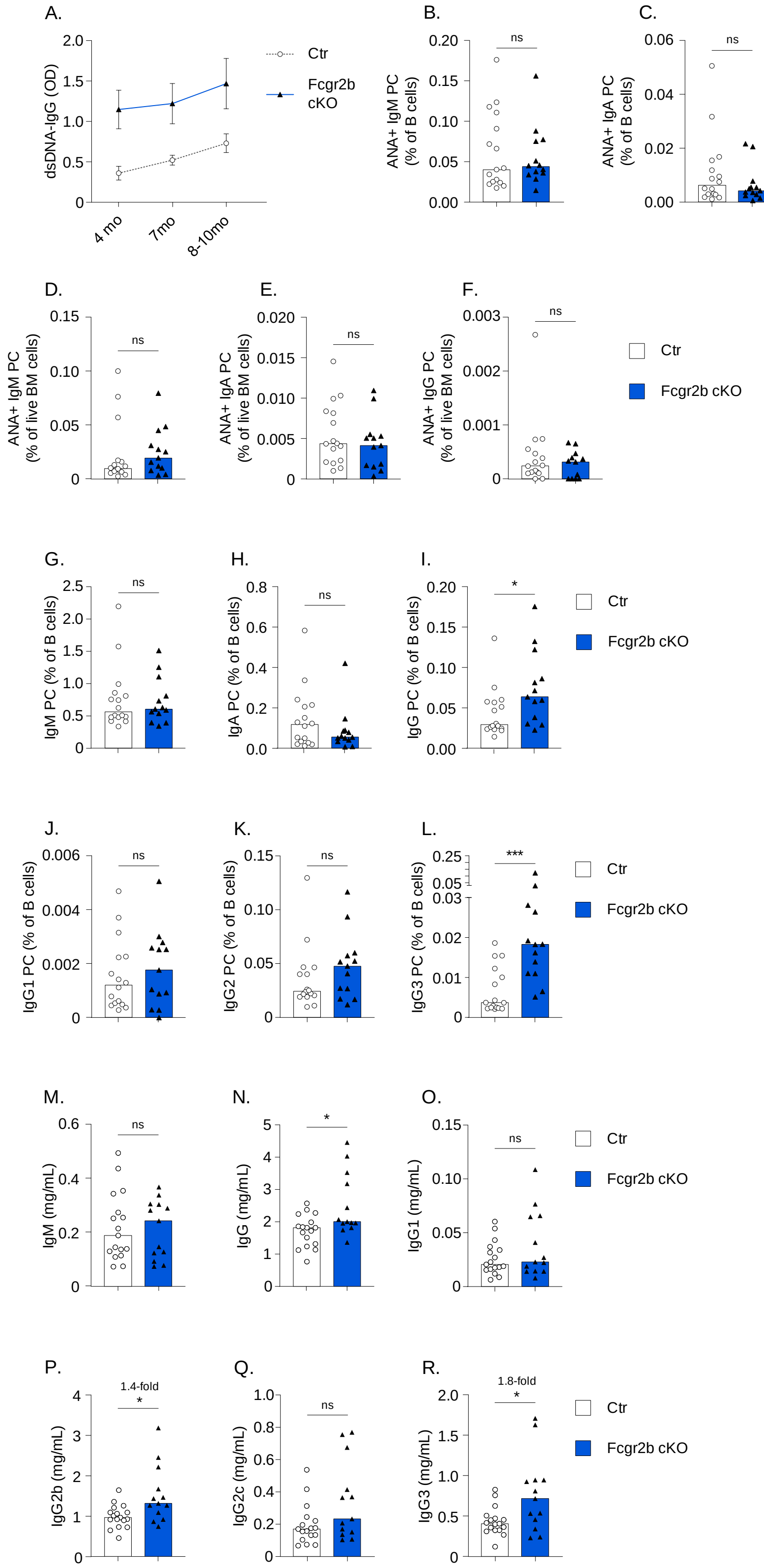


**Figure S1**: Detailed serum and PC analysis in *Fcgr2b* cKO mice

A) Timecourse of dsDNA IgG on serum of female Ctr and *Fcgr2b* cKO deficient mice. B-F) Frequency of ANA+ PCs in spleen and BM, by isotype. G-L) Frequency of PCs by isotype and IgG subclass, as a percent out of total B cells in spleen. M-R) Levels of total immunoglobulin, by isotype and IgG subclass, in serum of 10-12 month old female Ctr and *Fcgr2b* cKO mice.

Data are shown as mean +/- SEM (A) or median with each symbol representing an individual mouse (n=7 per group from 1 experiment in A; n=13-17 per group pooled from 3 independent experiments in other panels). Asterisks indicate significant differences (*p<0.05; **p<0.01; ***p<0.001) obtained using Mann whitney U test.

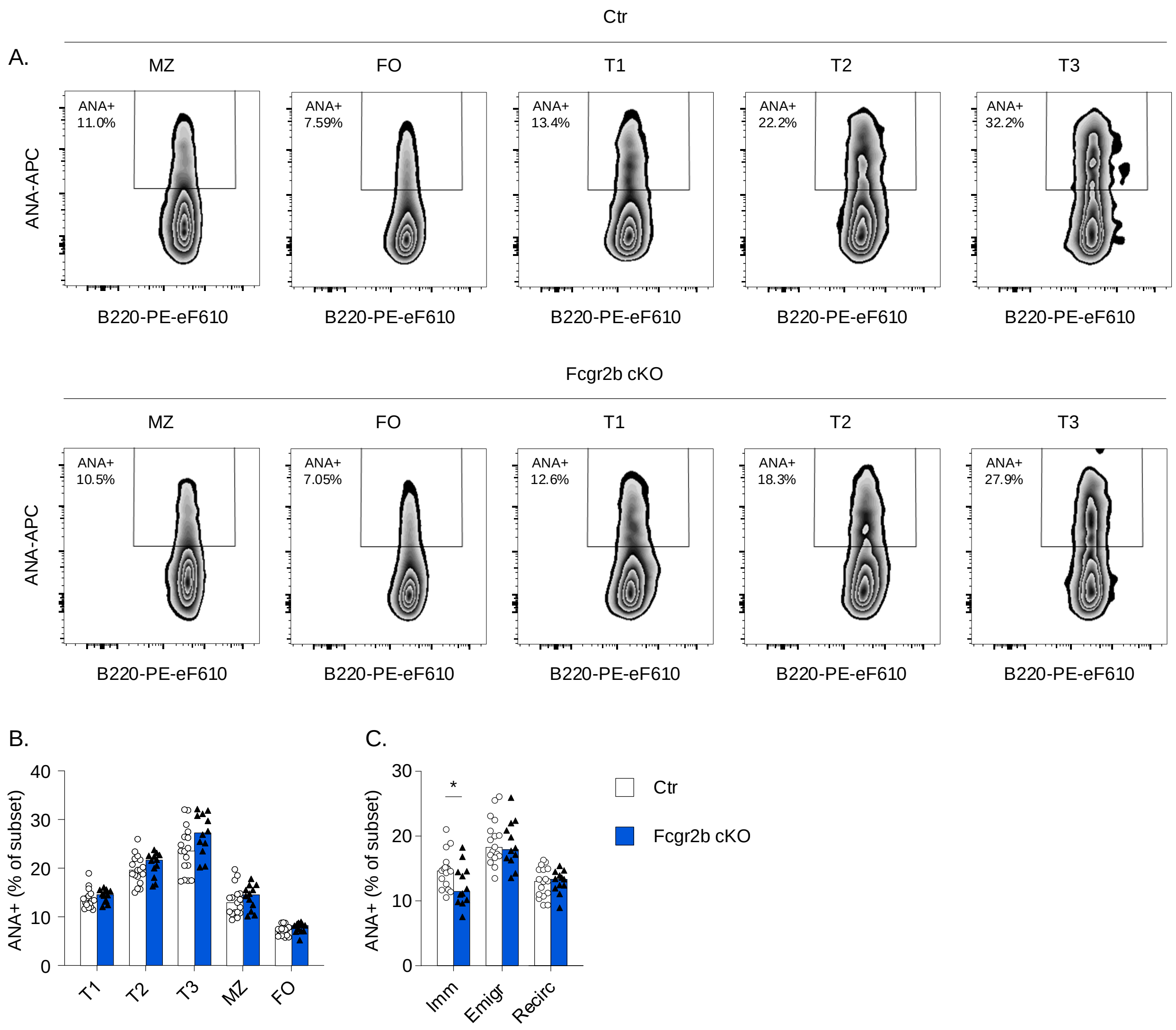


**Figure S2**: Early tolerance checkpoints for ANA in BM and spleen.

The frequency of autoreactive ANA+ B cells in 10-12 month old control (Ctr) and *Fcgr2b* cKO mice was established using flow cytometry through ANA surface staining. A) Representative example of ANA staining in B cell subsets in the spleen. B-C) Summary of ANA reactivity in B cell subsets in the spleen and bone marrow. B cell subsets were gated as described in Methods.

Data are shown as median with each symbol representing an individual mouse (n=12-17 per group pooled from 3 independent experiments). Asterisks indicate significant differences (*p<0.05; **p<0.01; ***p<0.001) obtained using Mann whitney U test.

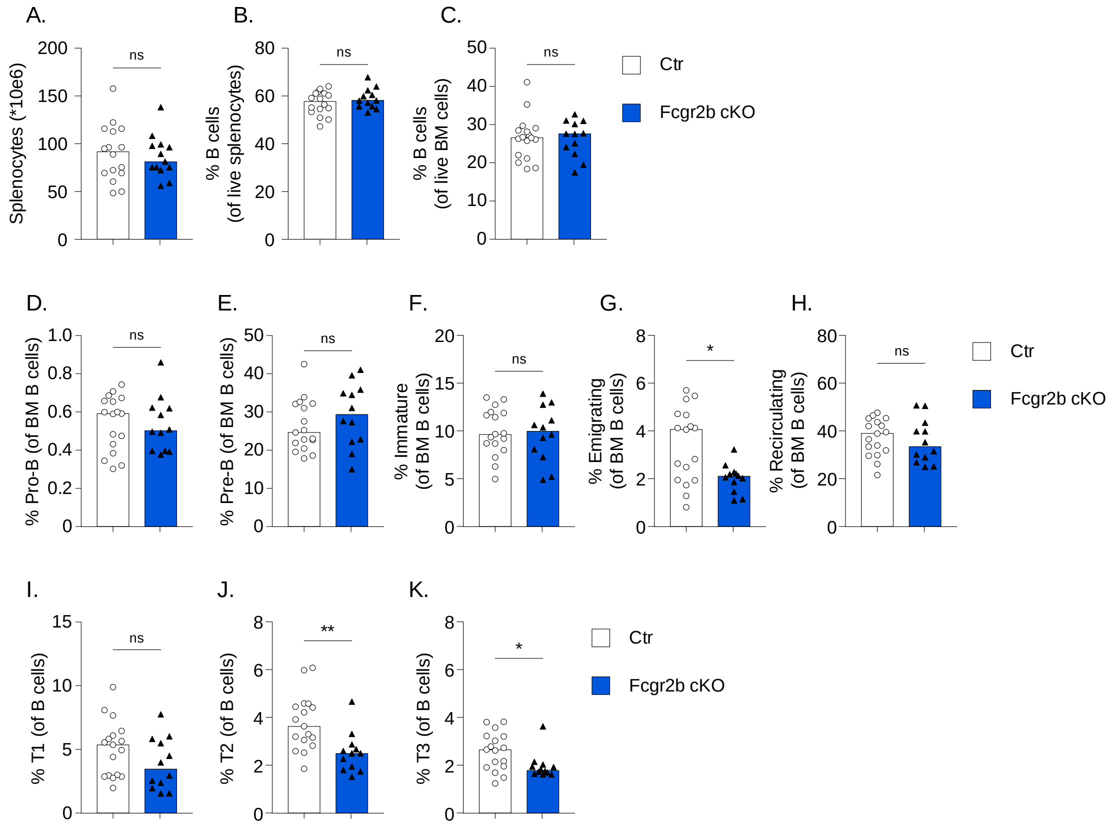


**Figure S3**: B cell subsets in BM and spleen.

The frequency of B cell subsets in spleen and bone marrow of 10-12 month old control (Ctr) and *Fcgr2b* cKO mice was established using flow cytometry. A) Total splenocyte count. B,C) Frequency of total B220+ B cells in spleen and bone marrow. Representative example of ANA staining in B cell subsets in the spleen. C-M) Summary of B cell subset frequencies in the spleen and bone marrow. B cell subsets were gated as described in Methods.

Data are shown as median with each symbol representing an individual mouse (n=12-17 per group pooled from 3 independent experiments). Asterisks indicate significant differences (*p<0.05; **p<0.01) obtained using Mann whitney U test.

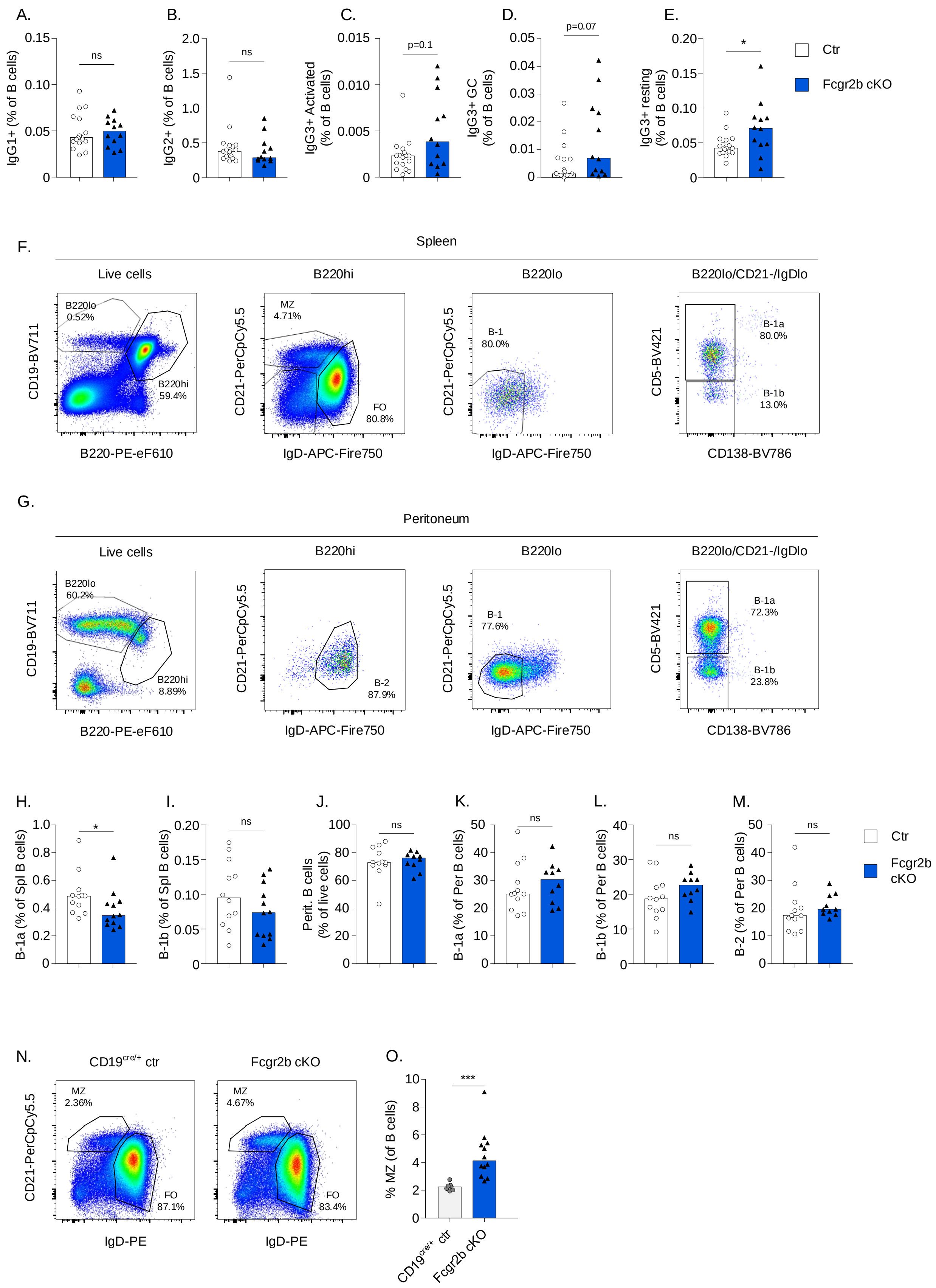
A.
B.
C.
D.
E.
IgG1+ (% of B cells)
IgG2+ (% of B cells)
IgG3+ Activated (% of B cells)
IgG3+ GC (% of B cells)
IgG3+ resting (% of B cells)
ns
ns
p=0.1
p=0.07
*
Ctr
Fcgr2b cKO
F.
Spleen
Live cells
B220hi
B220lo
B220lo/CD21-/IgDlo
B220lo 0.52%
B220hi 59.4%
MZ 4.71%
FO 80.8%
B-1 80.0%
B-1a 80.0%
B-1b 13.0%
CD19-BV711
CD21-PerCpCy5.5
CD5-BV421
B220-PE-eF610
IgD-APC-Fire750
CD138-BV786
G.
Peritoneum
B220lo 60.2%
B220hi 8.89%
B-2 87.9%
B-1 77.6%
B-1a 72.3%
B-1b 23.8%
H.
I.
J.
K.
L.
M.
B-1a (% of Spl B cells)
B-1b (% of Spl B cells)
Perit. B cells (% of live cells)
B-1a (% of Per B cells)
B-1b (% of Per B cells)
B-2 (% of Per B cells)
*
ns
N.
CD19cre/+ ctr
Fcgr2b cKO
MZ 2.36%
FO 87.1%
MZ 4.67%
FO 83.4%
CD21-PerCpCy5.5
IgD-PE
O.
% MZ (of B cells)
***

**Figure S4**: IgG class-switched cells, B-1 and MZ B cells in *Fcgr2b* cKO mice.

A,B) Frequency of IgG1+ and IgG2+ B cells in 10-12 month old female control (Ctr) and *Fcgr2b* cKO mice. C-E) Frequency of IgG3+ B cell subsets based on CD38 and GL7 staining. F,G) Representative examples of B cell staining strategies in spleen and peritoneum. B220hi = B-2 cells and B220o = B-1 cells, further confirmed as CD21loIgD- and separated into B-1a and B-1b based on CD5 (right panel). H-M) Frequency of B-1 cells in spleen (H,I) and total B cells, B-1a, B-1b, and B-2 cells in peritoneum (J-M). N,O) Representative example and summary of MZ frequency within mature CD93- B cells.

Data are shown as median with each symbol representing an individual mouse (n=12-17 per group for A-E; n=10-12 per group for H-M; n=8=12 per group for N-O; each pooled from 2-3 independent experiments). Asterisks indicate significant differences (*p<0.05; **p<0.01; ***p<0.001) obtained using Mann whitney U test.

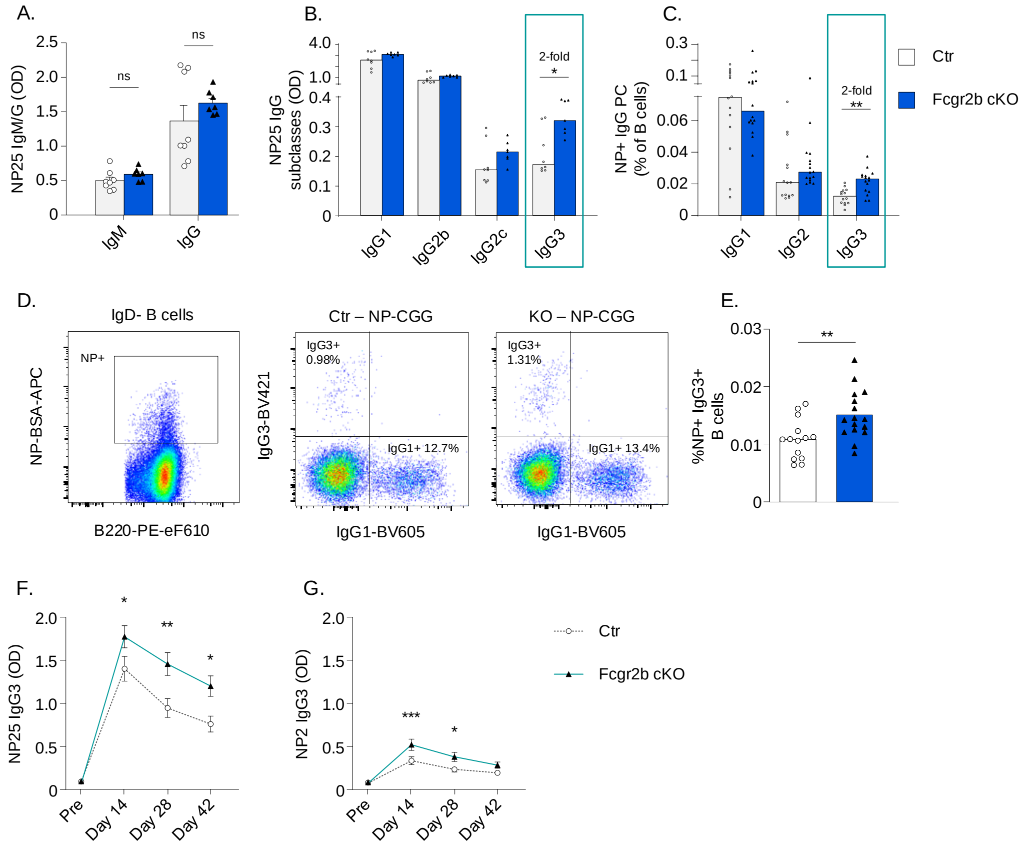


**Figure S5:** Characterization of T-dependent response to NP-CGG.

Female Ctr and *Fcgr2b* cKO mice were immunized with NP-CGG in Alum. Serum and splenocytes were obtained after 7 days (except F-H). A,B) Levels of NP-specific antibodies, separated by isotype and subclass. C) Frequency of NP-specific PCs in spleen as percentage of B cells, separated by IgG subclass. D) Representative example of surface NP gating on IgD- B cells (left) and IgG1 and IgG3 staining in IgD-NP+ B cells (middle and right). E) Frequency of IgG3+ NP+ B cells out of total B cells. F-G) Timecourse of serum titers for NP25- and NP2-specific IgG3.

Data are shown as median with each symbol representing an individual mouse (n=7-8 per group for A,B; n=14-16 for C,E; n = 17-20 for F-G; each pooled from 2-3 independent experiments). Asterisks indicate significant differences (*p<0.05; **p<0.01; ***p<0.001) obtained using Mann Whitney U test (A-E) or Two-way ANOVA with Bonferroni posthoc test (F-G).

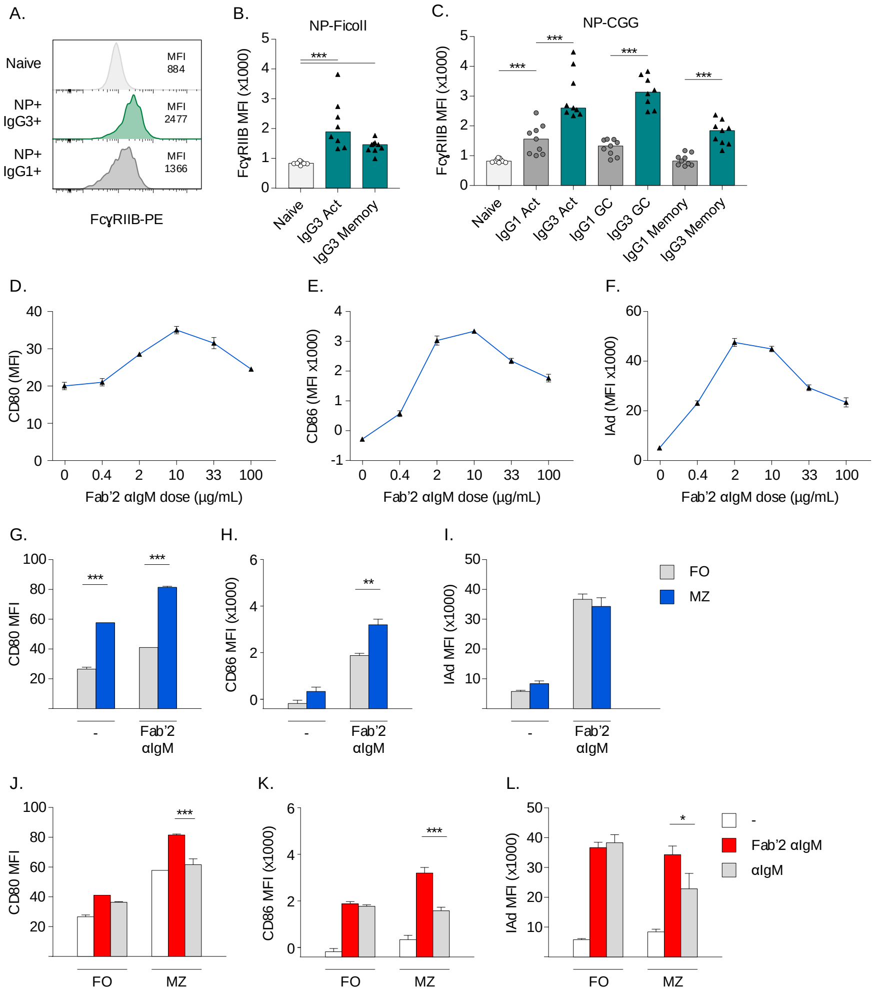


**Figure S6**: FcγRIIB expression and In vitro activation of follicular (FO) and marginal zone (MZ) B cells using intact and Fab'2 anti-IgM antibodies.

A) Representative example of staining for FcγRIIB in NP+ IgG1+ and IgG3+ B cells in control mice immunized with NP-CGG. B,C) FcγRIIB staining intensity in Naïve B cells, and different subsets of IgG1+ and IgG3+ B cells following immunization with NP-Ficoll and NP-CGG (day 7). D-L) FO and MZ B cells were sorted from the spleens of Ctr and *Fcgr2b* cKO mice, followed by stimulation with intact or equimolar concentrations of Fab'2 anti-IgM for 20 hours. Activation was measured by flow cytometry. D-F) Titration of Fab'2 anti-IgM stimulation in FO B cells. G-I) Comparison of activation of FO and MZ B cells in Ctr mice. J-L) Comparison of intact versus Fab'2 anti-IgM in FO and MZ B cells.

Data are shown as median (B,C) or mean +/- SEM (D-L) (n=8-9 per group for B-C; n=4-7 per group for G-L; each pooled from 2-3 independent experiments; except D-F which was from 1 experiment with 2 mice). Asterisks indicate significant differences (*p<0.05; **p<0.01; ***p<0.001) obtained using Mann Whitney U test (B,C) or Two-way ANOVA with Bonferroni posthoc test (G-L).

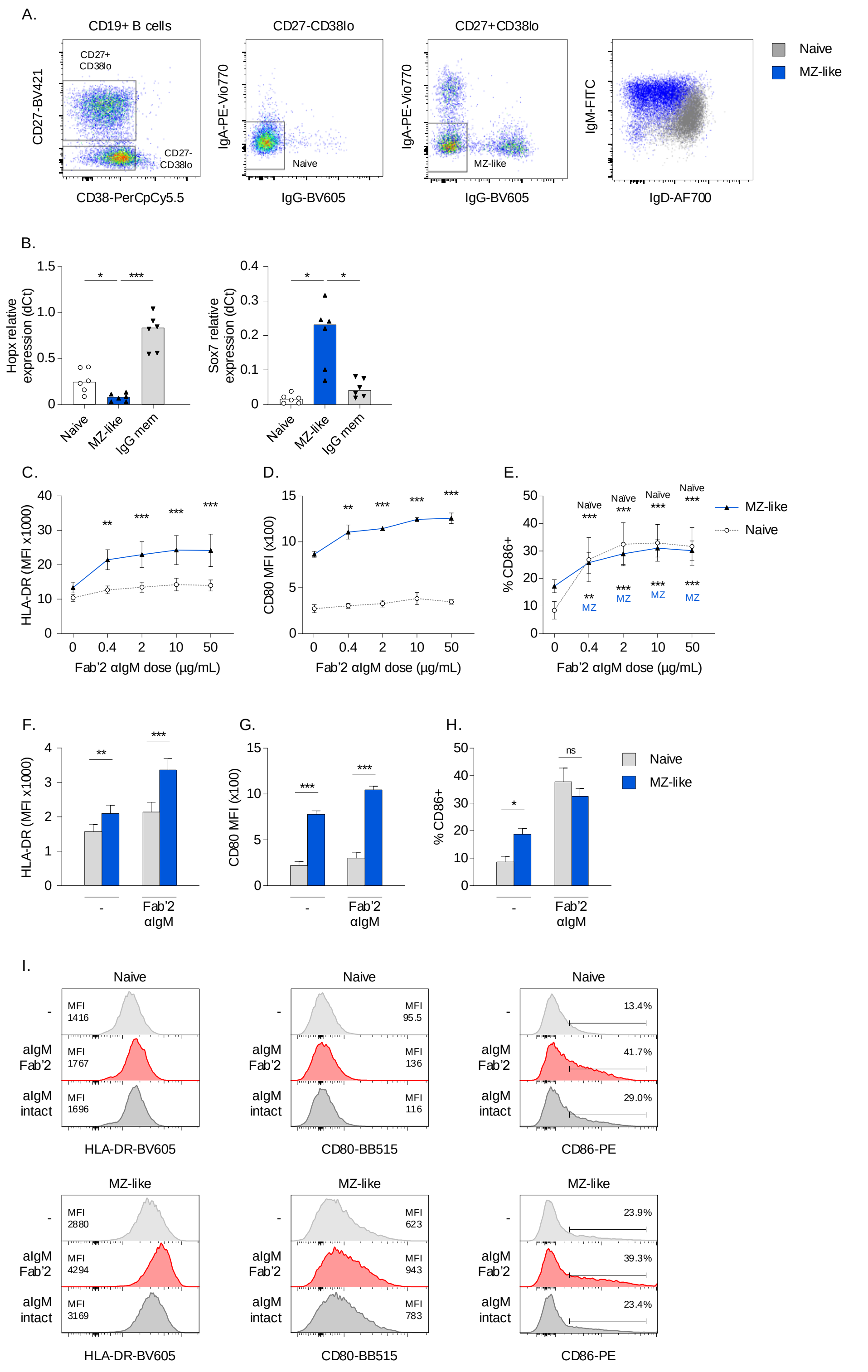


**Figure S7**: Human MZ-like B cell phenotype, titration of aIgM, and effect of Fab'2 anti-IgM on B cell activation, A) Gating strategy for human Naive and MZ-like B cells, starting with live CD19+ B cells. Cells were sorted from PBMCs of healthy donors, and analyzed by flow cytometry. B) qPCR for *HOPX* and *SOX7* in sorted B cell subsets, confirming MZ-like phenotype. Relative expression was normalized to polr2a. C-I) sorted Naïve and MZ-like B cells which were stimulated with intact anti-IgM or equimolar concentrations of Fab'2 anti-IgM 20 hours. Activation was measured by flow cytometry. C-E) Titration of Fab'2 anti-IgM in sorted Naïve and MZ-like B cells. F-H) Comparison of Naïve and MZ-like B cell activation, stimulated with 2 ug/mL Fab'2 anti-IgM. I) Representative examples of sorted Naïve and MZ-like B cells which were stimulated with 3 ug/mL intact anti-IgM or equimolar concentrations (2 ug/mL) of Fab'2 anti-IgM 20 hours. Activation was measured by flow cytometry.

Data are shown as median with each symbol representing an individual donor (B) or mean +/- SEM (C-H) (n=4-10 per group from 2-5 independent experiments for B; n=4 per group from 2 independent experiments for C-E; n=10 per group from 5 independent experiments for F-H). Asterisks indicate significant differences (*p<0.05; **p<0.01; ***p<0.001) obtained using Wilcoxon signed rank test (B) or Two-way ANOVA with Bonferroni posthoc test (C-H).

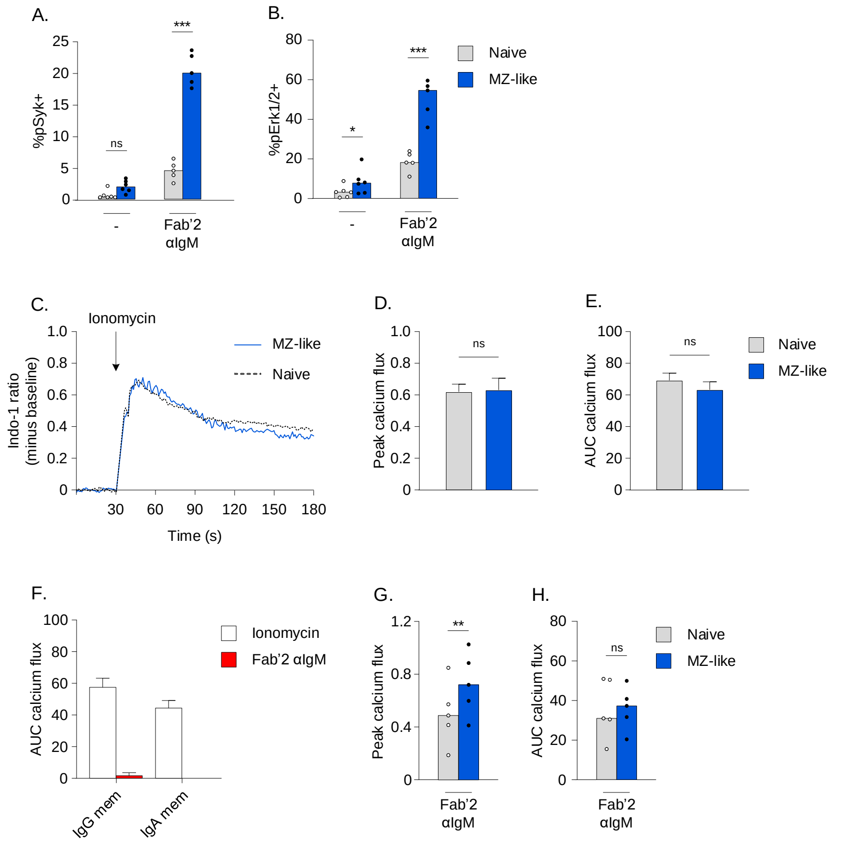


**Figure S8**: Phosphoflow and calcium flux with Fab'2 anti-IgM.

A,B) PBMCs from healthy donors were stimulated with 2 ug/mL of Fab'2 anti-IgM for 60 min, after which the phosphorylation of signaling molecules was analysed by phosphoflow. Comparison of phosphorylation of Syk and Erk in Naïve versus MZ-like B cells. C-H) PBMCs from healthy donors were labeled with Indo-1 for calcium flux measurements. After 30 seconds of baseline measurement, 75 ug/mL intact anti-IgM or equimolar concentrations (50 ug/mL) of Fab'2 anti-IgM were added and the measurement continued for 2.5 min. 1 ug/mL Ionomycin was used as a positive control. Peak calcium flux and area under the curve (AUC) were calculated using Flowjo. C-E) Representative examples and summary of Ionomycin-induced calcium flux in Naïve and MZ-like B cells gated as in

Figure S7A. F) Lack of calcium flux in IgG+ and IgA+ B cells following Fab’2 anti-IgM in comparison to Ionomycin as positive control. G,H) Comparison of the peak and AUC of calcium flux in Naïve and MZ-like B cells following stimulation with Fab’2 anti-IgM.

Data are shown as mean +/- SEM (n=5-6 per group from 2-3 independent experiments). Asterisks indicate significant differences (*p<0.05; **p<0.01; ***p<0.001; Ns: not significant) obtained using Two-way ANOVA with Bonferroni posthoc test (A,B) or Mann Whitney U test (D-H).

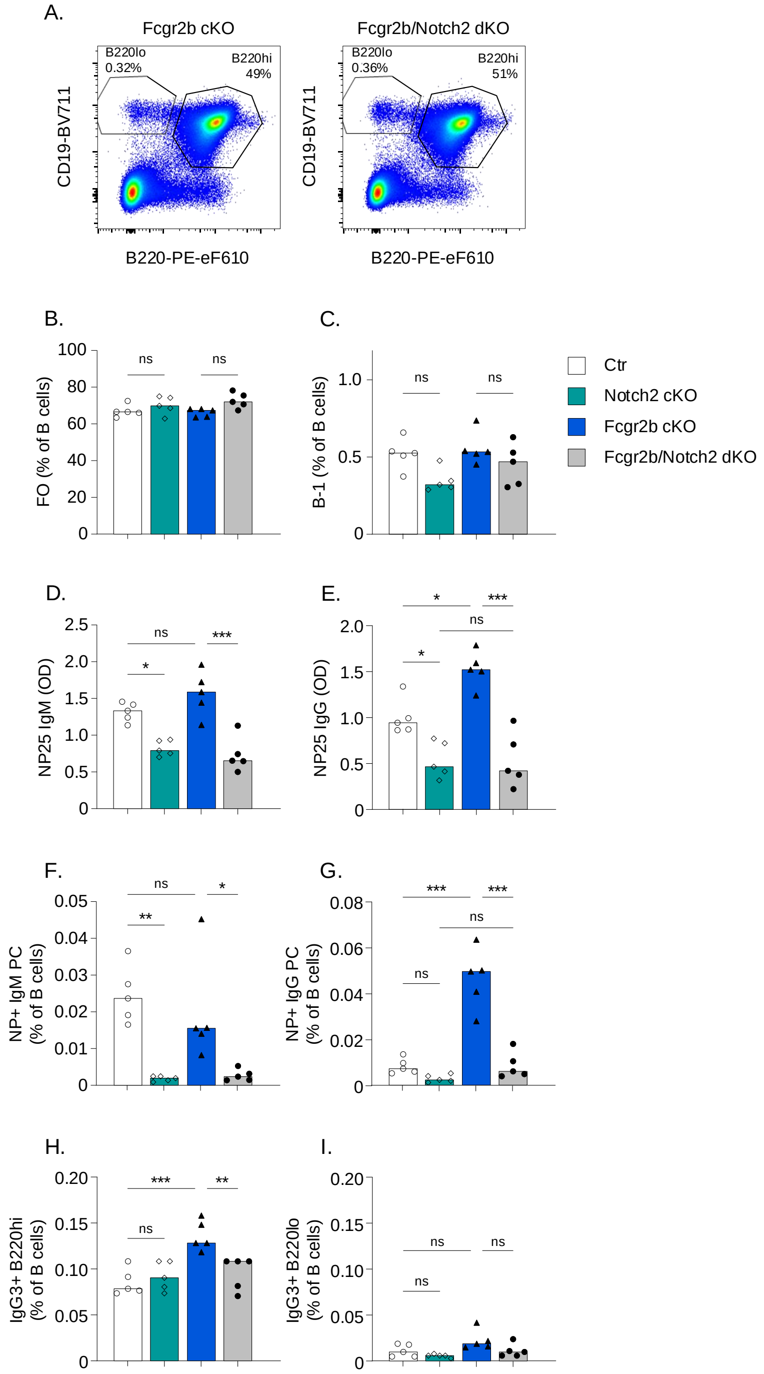


**Figure S9:** Combined FcɣRIIB and MZ deficiency reverses enhanced extrafollicular responses.

Female Ctr, *Notch2* cKO, *Fcgr2b* cKO, and *Notch2*/*Fcgr2b* dKO mice were immunized with NP-Ficoll. Serum and splenocytes were obtained after 7 days. A) Representative examples of gating for B220hi and B220lo. B,C) Frequencies of follicular (FO) and B-1 cells in spleen. D,E) Levels of NP-specific IgM and IgG in serum. F,G) Frequency of NP-specific IgM and IgG PCs in spleen. H,I) Frequency of IgG3+ B220hi (B-2) and IgG3+ B220lo (B-1) B cells in spleen.

Data are shown as median with each symbol representing an individual mouse (n=5 per group). Asterisks indicate significant differences (*p<0.05; **p<0.01; ***p<0.001) obtained using Two-way ANOVA with Bonferroni posthoc test.

**Table S1**: Flow cytometry antibodies

| Target species | Antigen | Fluorochrome | Clone | Company | Catalog Number |
|---|---|---|---|---|---|
| Mouse | B220 | PE-eF610 | RA3-6B2 | eBioscience | 61-0452-82 |
| Mouse | CD5 | BV421 | 53-7.3 | Biolegend | 100629 |
| Mouse | CD11c | AF700 | N418 | Biolegend | 117319 |
| Mouse | CD19 | BV711 | 6D5 | Biolegend | 115555 |
| Mouse | CD21 | PerCp-Cy5.5 | 7E9 | Biolegend | 123416 |
| Mouse | CD23 | BV786 | B3B4 | BD Biosciences | 563988 |
| Mouse | CD23 | FITC | B3B4 | Biolegend | 101605 |
| Mouse | CD23 | PECy7 | B3B4 | eBioscience | 25-0232-82 |
| Mouse | CD24 | PE-Cy7 | M1/69 | eBioscience | 25-0242-80 |
| Mouse | CD32b | APC | AT130-2 | eBioscience | 17-0321-82 |
| Mouse | CD32b | PE | AT130-2 | eBioscience | 12-0321-80 |
| Mouse | CD38 | BV711 | 90/CD38 | BD Biosciences | 740697 |
| Mouse | CD38 | PE | 90/CD38 | BD Biosciences | 553764 |
| Mouse | CD43 | FITC | S7 | BD Biosciences | 561856 |
| Mouse | CD80 | FITC | 16-10A1 | BD Biosciences | 561954 |
| Mouse | CD86 | AF700 | GL-1 | Biolegend | 105024 |
| Mouse | CD93 | BV421 | AA4.1 | BD Biosciences | 747716 |
| Mouse | CD93 | PE | AA4.1 | eBioscience | 12-5892-81 |
| Mouse | CD93 | PE-Cy7 | AA4.1 | BioLegend | 136506 |
| Mouse | CD95 | BV421 | Jo2 | BD Biosciences | 562633 |
| Mouse | CD95 | PE-Cy7 | Jo2 | BD Biosciences | 557653 |
| Mouse | CD138 | BV786 | 281-2 | BD Biosciences | 740880 |
| Mouse | GL7 | eF450 | GL7 | eBioscience | 48-5902-82 |
| Mouse | GL7 | FITC | GL7 | Biolegend | 144604 |
| Mouse | IAd | APC | AMS-32.1 | eBioscience | 17-5323-80 |
| Mouse | IgA | FITC | C10-3 | BD Biosciences | 559354 |
| Mouse | IgD | APC-Fire750 | 11-26c.2a | Biolegend | 405744 |
| Mouse | IgD | PE | 11-26C.1 | BD Biosciences | 558597 |
| Mouse | IgG1 | BV605 | A85-1 | BD Biosciences | 563285 |

| **Target species** | **Antigen** | **Fluorochrome** | **Clone** | **Company** | **Catalog Number** |
|---|---|---|---|---|---|
| Mouse | IgG1 | FITC | A85-1 | BD Biosciences | 553443 |
| Mouse | IgG2 | BV711 | R2-40 | BD Biosciences | 744296 |
| Mouse | IgG2 | BV786 | R2-40 | BD Biosciences | 744297 |
| Mouse | IgG3 | BV421 | R40-82 | BD Biosciences | 565808 |
| Mouse | IgG3 | BV605 | R40-82 | BD Biosciences | 744135 |
| Mouse | IgM | APC-eF780 | II/41 | eBioscience | 47-5790-82 |
| Mouse | IgM | eF450 | II/41 | eBioscience | 48-5790-82 |
| Mouse | IgM | PE | II/41 | eBioscience | 12-5790-81 |
| Mouse | IgM | PE-Cy7 | eB121-15F9 | eBioscience | 25-5890-82 |
| Mouse/ Human | Erk1/2 (pT202/pY204) | AF647 | 20A | BD Biosciences | 612593 |
| Mouse/ Human | Syk (pY348) | PE | I120-722 | BD Biosciences | 558529 |
| Human | CD3 | BV510 | UCHT1 | Biolegend | 300447 |
| Human | CD14 | BV510 | M5E2 | Biolegend | 301841 |
| Human | CD19 | PE-CF594 | HIB19 | BD Biosciences | 562321 |
| Human | CD19 | PerCp-eF710 | SJ25C1 | eBioscience | 46-0198-42 |
| Human | CD27 | BV421 | M-T271 | BD Biosciences | 562513 |
| Human | CD32 | PE | FLI8.26 | BD Biosciences | 550586 |
| Human | CD38 | BV711 | HIT2 | Biolegend | 303528 |
| Human | CD80 | BB515 | L307.4 | BD Biosciences | 565008 |
| Human | CD86 | PE | 2331 /FUN-1 | BD Biosciences | 555658 |
| Human | HLA-DR | BV605 | L243 | Biolegend | 307640 |
| Human | IgA | PE-Vio770 | IS11-8E10 | Miltenyi | 130-114-003 |
| Human | IgD | AF700 | IA6-2 | Biolegend | 348229 |
| Human | IgD | PE-CF549 | IA6-2 | BD Biosciences | 562540 |
| Human | IgG | BV605 | G18-145 | BD Biosciences | 563246 |
| Human | IgG | BV785 | G18-145 | BD Biosciences | 564230 |
| Human | IgM | FITC | MHM-88 | Biolegend | 314506 |
| | Mouse IgG1,k isotype control | AF647 | MOPC-21 | Biolegend | 400130 |

| **Target species** | **Antigen** | **Fluorochrome** | **Clone** | **Company** | **Catalog Number** |
|---|---|---|---|---|---|
| | Mouse IgG1,k isotype control | BB515 | X40 | BD Biosciences | 564416 |
| | Mouse IgG1,k isotype control | PE | (P3.6.2.8.1 | eBioscience | 12-4714-82 |
| | Mouse IgG2a,k isotype control | APC | eBM2a | eBioscience | 17-4724-81 |
| | Mouse IgG2a,k isotype control | BV605 | G155-178 | BD Biosciences | 562778 |
